# Structural Identification of Major Molecular Determinants for Phosphotyrosine Recognition in Tyrosine Kinases Reveals Tumour Promoting and Suppressive Functions

**DOI:** 10.1101/2025.06.10.658871

**Authors:** Nuo Cheng, Luis R. Millán-Barea, Marc Creixell, Alexis R. Barr, Yi Wen Kong, Brian A. Joughin, Cécile K. Lopez, Jette Lengefeld, James Clarke, Chuan Liu, Ganapathy Sriram, Tania J. González-Robles, Bert van de Kooij, Simonas Savickas, Erwin M. Schoof, Jared L. Johnson, Chris Bakal, Lewis C. Cantley, Roberto Chiarle, Justin Pritchard, Stevan Hubbard, Brian J.P. Huntly, Michael Hemann, Pau Creixell, Michael B. Yaffe

**Affiliations:** Cancer Research UK Cambridge Institute, University of Cambridge, UK; Center for Precision Cancer Medicine, Koch Institute for Integrative Cancer Research and Departments of Biological Engineering and Biology, Massachusetts Institute of Technology, Cambridge MA, USA; MRC Laboratory of Medical Sciences, London, UK; Institute of Clinical Sciences, Faculty of Medicine, Imperial College London, London, UK; Department of Haematology, University of Cambridge, Cambridge, UK; Cambridge Stem Cell Institute, Cambridge, UK; Helsinki Institute of Life Science, HiLIFE, Institute of Biotechnology, Faculty of Biological and Environmental Sciences, University of Helsinki, Helsinki, Finland; Center for Hematology and Regenerative Medicine, Department of Medicine Huddinge, Karolinska Institutet, Stockholm, Sweden; Department of Biomedical Engineering, Pennsylvania State University, University Park, State College, PA, 16802, USA; Department of Biochemistry and Molecular Pharmacology, New York University Grossman School of Medicine, New York, New York, USA; Department of Biotechnology and Biomedicine, Technical University of Denmark, Søltofts Plads B221, Kgs. Lyngby, 2800, Denmark; Meyer Cancer Center, Weill Cornell Medicine, New York, NY 10021, USA; Department of Cell Biology, Harvard Medical School, Boston, MA 02115, USA; Dana-Farber Cancer Institute, Harvard Medical School, Boston, MA 02215, USA; Dynamical Cell Systems Group, Division of Cancer Biology, Institute of Cancer Research, 237 Fulham Road, London, UK; Department of Pathology, Boston Children’s Hospital and Harvard Medical School, Boston, MA 02115, USA; Department of Molecular Biotechnology and Health Sciences, University of Torino, 10126 Torino, Italy; Division of Hematopathology, IEO European Institute of Oncology IRCCS, Milan, Italy

## Abstract

Protein tyrosine kinases activate signaling pathways by catalyzing the phosphorylation of tyrosine residues in their substrates. Mounting evidence suggests that, in addition to recognizing phosphorylated tyrosine (pTyr) residues through specific phosphobinding modules, many protein kinases selectively recognize pTyr directly adjacent to the tyrosine residue they phosphorylate and catalyze the formation of twin pTyr-pTyr sites. Here, we demonstrate the importance of this phosphopriming-driven twin pTyr signaling in promoting cell cycle progression through the cell cycle-inhibitory protein p27^Kip1^. We identify, structurally resolve, and tune two distinct molecular determinants driving the selective recognition of pTyr directly N- and C-terminal to the target phospho-acceptor tyrosine site. We further show structural and biochemical conservation in this recognition, and identify cancer-associated alterations to these determinants that are unable to recognize phosphoprimed substrates. Finally, using an *in vivo* mouse model of leukemia we show that Bcr-Abl mutants unable to recognize phosphoprimed substrates paradoxically result in enhanced tumor development and progression. These data indicate that Bcr-Abl, like other proto-oncogenes such as Ras or Myc, engages both pro- and anti-oncogenic programs – but in the case of Bcr-Abl, this is accomplished through a mechanism involving traditional and phosphoprimed substrate recognition.

## Introduction

Cells respond to internal and external cues by conditionally activating signaling pathways and downstream molecular programs. The conditional activation of these pathways most commonly occurs through the post-translational modifications of proteins which are subsequently specifically recognized by downstream signaling components, ensuring precise and temporally controlled signal transduction. In phosphorylation-based signaling systems, for instance, upon becoming active, protein tyrosine kinases act as “writers” by phosphorylating tyrosine residues in their protein substrates^1–4^ to generate phosphorylated tyrosine (henceforth referred to as pTyr). These pTyr sites are subsequently recognized and bound by phospho-binding domains (“readers”), such as SH2 (Src Homology 2) and PTB (pTyr-binding) domains^5^, or by protein tyrosine phosphatases that remove phosphorylation, reverting them back to tyrosine sites (“erasers”)^6^.

The discovery of specific protein domains capable of independently “writing”, “reading” or “erasing” these post-translational modifications has provided a basis to consider signaling systems as highly modular^7, 8^. Despite this modularity, it is becoming increasingly clear that, far from being independent steps, many of these processes co-occur and are tightly interrelated. Initial evidence for this interrelation emerged from the observation that “reader” and “writer” protein domains often co-occur within the same protein, for instance in protein tyrosine kinases containing SH2 domains^9, 10^. Harboring “reader” domains enables protein kinases to recruit previously phosphorylated tyrosine substrates (a process often referred to as “priming” phosphorylation) and phosphorylate them at additional sites^11–16^. In contrast to pTyr signaling utilizing independent “reader” and “writer” domains with spatially separated pTyr sites to propagate downstream signaling events, we and others have more recently discovered that “priming” phosphorylation events can also occur at positions directly adjacent to each other^17–20^.

Notably, recognition of pTyr-Tyr (pYY) or Tyr-pTyr (YpY) sites and their subsequent phosphorylation to form twin pTyr-pTyr (pYpY) sites can only be possible if the protein kinase domain itself can recognize pTyr residues. In other words, the recognition of pTyr by the kinase domain is an indispensable prerequisite for “priming” signaling and, in doing so, some protein kinase domains effectively function as both “readers” and “writers”. Given this, two fundamental biological questions remain unresolved: how does pTyr recognition take place at a structural and molecular level, and wWhat effects do perturbations in these systems have in normal and aberrant cell signaling, such as in cancer?

Here, we show that a large subset of protein tyrosine kinases is capable of selectively recognizing pTyr directly adjacent to the tyrosine residue they phosphorylate. Initially focusing on the well-studied non-receptor tyrosine kinases Abl and Src, that are both aberrantly activated in cancer, our data shows that while Abl selectively recognizes pTyr directly N-terminal to its target tyrosine, Src conversely recognizes pTyr directly C-terminal to the target tyrosine. Using Abl and Src as exemplars, we identify the two distinct molecular and structural determinants driving the selective recognition of pTyr directly N-terminal and C-terminal to their target phospho-acceptor tyrosine site. By combining biochemical screening of other non-receptor and receptor tyrosine kinases and solving the crystal structure of the insulin receptor kinase (IRK), we expand our understanding of how tyrosine kinases recognize pTyr.

Finally, to evaluate its cellular relevance, we disrupt twin pTyr signaling in two different ways. First, we perturb the shared Abl and Src substrate p27^Kip1^ and show that phosphorylation of twin tyrosine (YY) sites positively promotes cell cycle progression. Second, by using structure-based mutants in an *in vivo* mouse model of leukemia, we show that Bcr-Abl variants with disrupted pTyr recognition led to more aggressive cancer progression, revealing a new anti-oncogenic signaling arm downstream of the pro-oncogenic Bcr-Abl kinase. While other pro-oncogenes like Ras or Myc are known to engage anti-oncogenic programs through collateral signaling pathways, our data suggest that Bcr-Abl can directly engage anti-oncogenic programs through its twin pTyr signaling arm.

## Results

### The Non-Receptor Tyrosine Kinases Abl and Src Conditionally Phosphorylate Twin Tyrosine Substrates by Selectively Recognizing pTyr

Recent results from our lab and others^17, 19, 20^ have revealed, using position-specific peptide libraries (PSPL, Figure S1A), that a surprising number of tyrosine kinases tolerate, or even prefer, a previously phosphorylated amino acid in one of the sites proximal to the tyrosine phosphoacceptor residue. These tyrosine kinases can thereby catalyze the creation of pYpY sites in their substrates.

Analysis of the human proteome (UP000005640)^21^ revealed 28,637 twin YY sites, of which 1,746 have been annotated as being phosphorylated on at least one of the two residues^22^. Proteins containing these YpY and pYY sites were primarily cytosolic and were enriched in the Gene Ontology (GO) terms “protein binding”, “transmembrane receptor protein tyrosine kinase signaling pathway”, “kinase activity”, “protein autophosphorylation”, “protein-containing complex”, “protein autophosphorylation”, and “intracellular signal transduction” (Figure 1A) relative to the total proteome.

**Figure 1.**
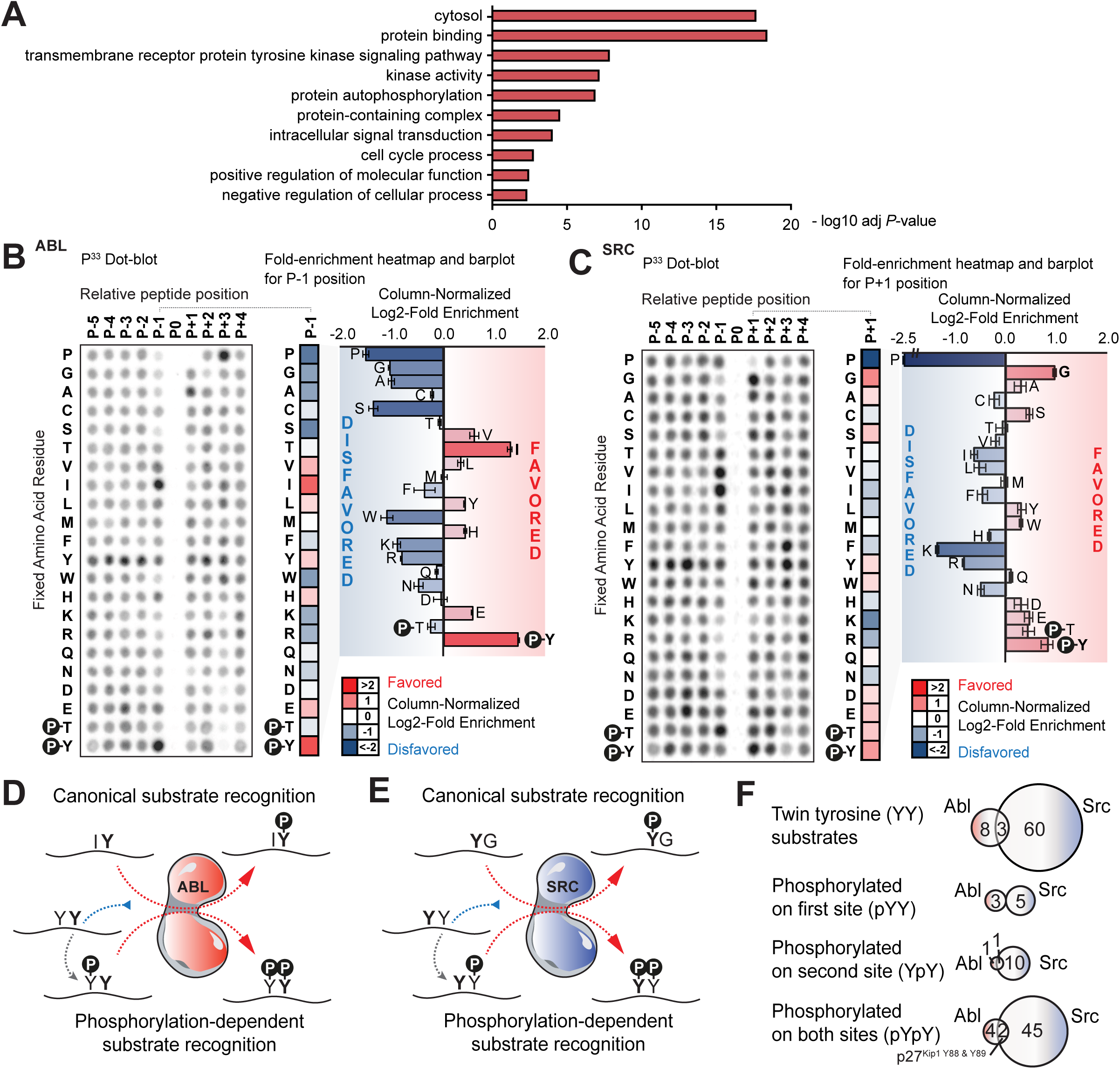
The Non-Receptor Tyrosine Kinases Abl and Src Kinases Conditionally Phosphorylate Twin Tyrosine Substrates by Selectively Recognizing pTyr. (A) Selected gene sets that are statistically significantly enriched in YY sites reported to be phosphorylated relative to all YY sites. (B) Positional Scanning Peptide Library (PSPL) screening results for Abl kinase shown on the left with the different fixed amino acid residues as rows (x axis) and different relative positions where the residues are fixed as columns (y axis). A column-normalized fold-enrichment heatmap for the relative position directly N-terminal from the phosphorylation site position (P-1) is shown in the middle with red denoting a favored amino-acid residue for Abl and blue denoting a disfavored one. To facilitate direct comparison between residues and their enrichment, the same fold-enrichment values are displayed as barplots. (C) Positional Scanning Peptide Library (PSPL) screening results for Src kinase shown as in B, but with a focus on the P+1 position. (D, E) Graphical representation of the canonical and non-canonical, phosphorylation-dependent motifs for Abl (D) and Src (E) kinases based on the data from panels B & C. (F) Top. Venn diagrams displaying phosphorylation sites reported in PhosphositePlus as being downstream of Abl kinase, Src kinase or both and that contain a tyrosine directly adjacent to them. Bottom. Further classification of these YY sites with those reported to be phosphorylated on the first N-terminal tyrosine, the second C-terminal or both. tyrosine residues 88 and 89 of p27^Kip1^ are highlighted as the only sites shared by Abl and Src that are reported to be phosphorylated on both residues (even if not necessarily simultaneously). See also Figure S1.

The four possible phosphorylation states of a tandem tyrosine motif (YY, pYY, YpY, pYpY) suggest a potentially broader regulatory set of mechanisms for tyrosine kinase signaling. First, the two individual sites might operate redundantly, with either phosphorylation being sufficient to enable a biological consequence (i.e., an “OR” gate). Second, they might operate to amplify signaling, with double phosphorylation resulting in a stronger consequence than either single phosphorylation alone (i.e., like an “AND” gate). Finally, the sites might operate combinatorially, with up to four totally independent consequences for the four possible phosphorylation states. Each of these possibilities allows the potential convergence of tyrosine kinase signaling pathways on YY-containing motifs in a substrate. To examine this, we focused on the first two oncogenic non-receptor tyrosine kinases ever identified, Src^23, 24^ and Abl^25, 26^, as examples of well-studied and clinically relevant model tyrosine kinases to study these functions.

Multiple independent experiments using the isolated kinase domains demonstrate that Abl strongly selects for pY in the P-1 position relative to the phospho-acceptor tyrosine site in its optimal substrate motif (Figures 1B and S1B), whereas, conversely, Src selects for pY in the P+1 position (Figures 1C and S1C). These mirroring pY motif preferences suggest that both tyrosine kinases could phosphorylate a subset of their substrates in a phosphopriming-dependent manner (Figures 1D-F), such that each kinase, in phosphorylating an unphosphorylated substrate, could produce a phosphoprimed substrate optimized for phosphorylation by the other kinase.

### Tyrosine Residues p27^Y88^ and p27^Y89^ Both Mediate Cell Cycle Progression in a Non-redundant Manner

Among all substrates of Abl and Src annotated in the resource PhosphositePlus^22^, we identified a single instance of consecutive tyrosine residues known to be phosphorylated by both of these kinases – Y88 and Y89 on the cell cycle inhibitor p27^Kip1^ (Figure 1F). To determine whether these two tandem phosphosites can both be present simultaneously in a single p27 molecule, we performed targeted mass spectrometry in lysates from cycling hTert-RPE1 cells. Figures 2A and S2A show the resulting mass spectrum, consistent with the expected mass-to-charge ratio for a twin-phosphorylated peptide, indicating that p27 protein in these freely cycling cells exists in the doubly phosphorylated form.

**Figure 2.**
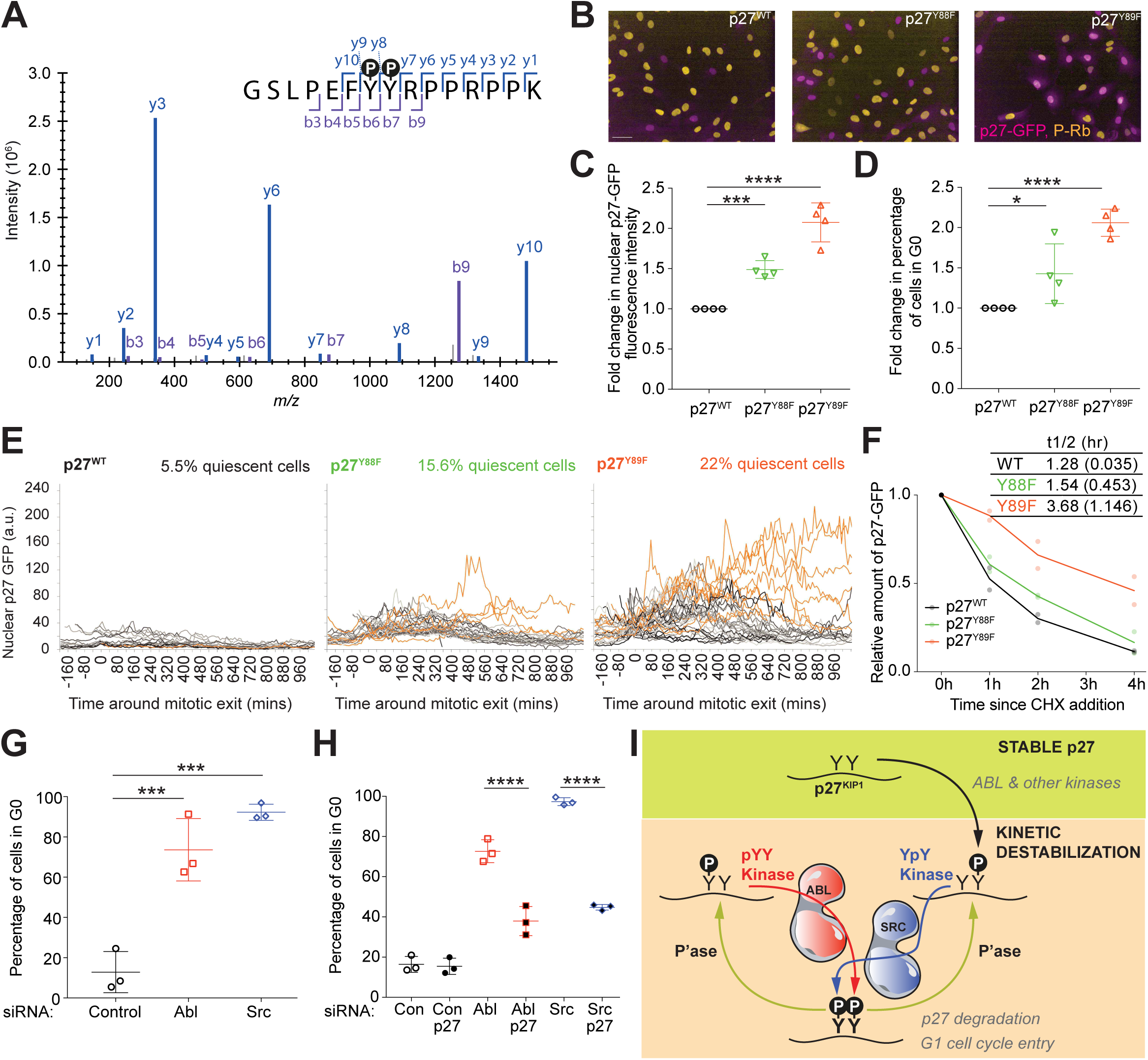
Phosphorylation of Tyrosine Residues p27^Y88^ and p27^Y89^ Is Required for Cell Cycle Progression. (A) Mass spectrum displaying intensity (y axis) as a function of mass-to-charge ratio (x axis) for the different fragment ions also displayed as b- and y-ions within the p27 peptide sequence containing the twin pTyr-pTyr sites (p27^pY88 pY89^). (B) Representative merged images of p27-GFP (magenta) in genotypically CRISPR-tagged p27^WT^, p27^Y88F^ and p27^Y89F^ cell lines, fixed and immuno-stained for pS807/811 Rb (gold). Scale bar is 50 μm. (C, D) Quantification of the fold-change in (C) nuclear p27-GFP fluorescence intensity and (D) percentage of quiescent (G0) cells in the mutant p27 cell lines relative to the p27 wild type cell line. Quiescence was scored as pSer807/Ser811-Rb negative. Mean +/- STD is shown of n=4. One-way ANOVA (*** *P* <0.0005, **** *P* <0.0001). (E) Graphs showing quantification of nuclear p27-GFP in individual cells from timelapse imaging of p27^WT^, p27^Y88F^ and p27^Y89F^ cell lines, aligned *in silico* to time of mitotic exit. Orange traces are cells that enter spontaneous quiescence (e.g., do not enter S-phase for at least 600 mins). Numbers in orange are the percentage of cells in each cell line that enter spontaneous quiescence. Number of cells per cell line: p27^WT^ n=18, p27^Y88F^ n=32, p27^Y89F^ n=41 cells. (F) Quantification of p27-GFP levels from Western blotting after treating cells with 100 μg/mL cycloheximide from panel Figure S2H, normalized to t=0h controls. Line represents mean of n=2, and table shows half-lives of p27^WT^, p27^Y88F^ and p27^Y89F^ proteins in hrs (+/-STD shown in brackets). (G) Quantification of the percentage of quiescent (G0, pRb-) cells after siRNA-based knock-down of Abl and Src compared to non-targeting control siRNA treatment. Graph shows the mean +/- STD of n=3. One-way ANOVA (*** *P* <0.0005). (H) Quantification of the percentage of quiescent cells (G0) after siRNA-based knock- down of p27 alone and in combination with Abl, Src or non-targeting control siRNA treatment. Data shown is mean +/- STD of n=3. One-way ANOVA (**** *P* <0.0001). (I) Model illustrating how different non-redundant phosphorylating events on p27 at Y88 and/or Y89 by Abl and Src could differentially regulate p27 degradation and G1 cell cycle entry. See also Figure S2.

Phosphorylation of p27 has been shown to inactivate p27 as a CDK inhibitor, and to enhance its subsequent ubiquitination and proteasomal degradation, resulting in cell progression from G1 into S-phase^27–29^. The Y88 and Y89 phosphorylation sites on p27 have been previously reported to have some level of redundancy in this regard^28–32^. Given the existence of the doubly phosphorylated form, and motif specificities compatible with sequential phosphorylation in either order, we wondered whether both p27^Y88^ and p27^Y89^ sites, as well as both Abl and Src, are necessary to optimally regulate the cell-cycle inhibitory functions of p27, compatible with phosphopriming at these residues.

Previous studies relied on overexpression of p27 mutants to analyse the functions of the Y88 and Y89 residues. Since overexpressing cell cycle inhibitors can perturb cell cycle progression, we wanted to examine the impact of Y88 and Y89 mutation on endogenous p27 levels and cell cycle dynamics in order to preserve stoichiometry between enzyme-substrate reactions. We therefore used hTert-RPE1 mRuby-PCNA cells, an established non-transformed, diploid cellular model system where endogenous PCNA has been labelled with mRuby, enabling the study and quantification of cell cycle progression and phenotypes^33^. To track p27 dynamics and cell cycle progression in real-time, we generated an hTert-RPE1 mRuby-PCNA cell line where we labelled endogenous p27 at the C-terminus with eGFP, using CRISPR/Cas9 gene-editing (Figures S2B). We have previously shown that p27-GFP replicates p27 dynamics in HeLa cells^34^ and, here we confirmed that addition of eGFP to endogenous p27 in hTert-RPE1 mRuby-PCNA cells did not affect cell growth, p27 localisation or the ability of p27 to bind CDK complexes (Figures S2C-F). CRISPR/Cas9 gene editing was then used to mutate the endogenous *CDKN1B* locus (encoding p27, tagged with GFP) to generate nonphosphorylatable phenylalanine (F) mutations at the Y88 or Y89 sites. Cell cycle progression of p27^WT^ or p27^mut^ cells was then analyzed by fixed and live-cell imaging (Figure 2B and Figure S2E). Unlike what was previously observed in NB4 leukaemia cells^30^, neither of the p27 mutants showed overt differences in either their ability to bind to Cdk2 and Cdk4, or in their nuclear localization relative to p27^WT^, suggesting that loss of tyrosine phosphorylation at either of these sites does not affect subcellular localisation of endogenous p27 in these epithelial cells (Figures S2E-F). Compared to p27^WT^-GFP cells, both the Y89 mutant and, to a lesser degree, the Y88 mutant demonstrated a significant increase in nuclear p27 intensity (Figures 2B-C, S2G) coupled with an increase in the fraction of cells entering quiescence (scored as phospho-Rb negative cells) under otherwise optimal (10% FBS) growth conditions (Figure 2D).

Live cell imaging and quantification of p27-GFP levels in individual asynchronously cycling cells were synchronized *in silico* relative to each cell’s mitotic exit, allowing comparison of p27^WT^ and p27^mut^ cells with respect to p27-GFP nuclear fluorescence intensity and associated cell cycle progression. In p27^WT^ cells, p27-GFP nuclear fluorescence remained low, with 5.5% cells remaining in G0/quiescence after mitosis. In contrast, p27-GFP nuclear fluorescence was higher in both the Y88F and Y89F p27^mut^ cells, resulting in a roughly three-fold or four-fold increase, respectively, in the percentage of cells remaining in G0/quiescence after mitosis (Figure 2E).

Since tyrosine phosphorylation of p27^Y88^ should promote CyclinA/CDK2-mediated phosphorylation of p27^T187^ and subsequent recognition of p27 by the SCF^Skp2^ complex followed by proteasomal degradation^27, 28^, we evaluated whether the increased p27 levels in p27^mut^-GFP cells were driven by an increased p27 protein stability. To confirm that the increased p27 levels in p27^mut^-GFP cells were driven by an increase in protein stability, protein synthesis was inhibited using cycloheximide and p27 stability evaluated by Western blotting cell lysates as a function of time (Figure S2H). Both Y88F and Y89F mutations prolong the half-life of p27, with the Y89F mutation displaying a particularly pronounced effect (Figure 2F). These findings support non-redundant roles for these two phosphorylation sites in mediating the stability of endogenous p27 and cell cycle entry in cells.

To clarify the roles of Abl and Src in p27-mediated cell cycle progression in hTert-RPE1 cells, we used both small molecule inhibitors and siRNAs to perturb kinase function. Treatment of hTert-RPE1 mRuby-PCNA p27^WT^-GFP cells with either Abl (Nilotinib) or Src (Saracatinib) inhibitors resulted in cell cycle arrest (Supplementary Movies 2A-C) with a larger effect seen upon Saracatinib treatment. These findings were confirmed using siRNA knockdown. Depletion of Abl or Src led to an increased fraction of cells entering quiescence (G0) (Figure 2G and Figure S2I). Notably, cell cycle arrest upon Abl or Src depletion was at least partially p27-dependent, since co-depletion of p27 with Abl or Src reduced the fraction of cells arresting in G0 (Figures 2H). As was seen with drug treatment, depletion of Src had a more pronounced effect than depletion of Abl. These data suggest that both Abl and Src promote cell cycle entry through the destabilisation of p27.

Phosphorylation of Y89 (a known Abl site^32, 35^) appears to be more important than Y88 for p27 destabilization and cell cycle entry. Tyr 89 is a better sequence match to the optimal substrate motif of Abl than that of Src, particularly if Y88 is already phosphorylated (Figures 1B and 1D). In contrast, Y88 is a better optimal substrate match for Src, especially in the context of phosphorylated Y89. Paradoxically however, Src activity appears to be more essential than Abl activity in destabilizing p27 and promoting S phase entry. One possible model capable of explaining these results is shown in Figure 2I. In this model, p27 with phosphorylation at either or both of the two tyrosine sites promotes p27 degradation, leading to cell cycle entry. We have previously shown that Y89 is more accessible than Y88 when both sites are non-phosphorylated^35^ suggesting that Y89 could be the site first phosphorylated by Abl, or another kinase. This creates a phosphoprimed substrate suitable for phosphorylation at Y88 by Src. Dephosphorylation of either the Y88 or Y89 site by phosphatases such as SHP2^36^ maintains a phosphoprimed substrate suitable for rapid rephosphorylation by a kinase with substrate specificity for -1 or +1 pY, respectively. This would oppose complete dephosphorylation of p27 at Y88 and Y89, and effectively kinetically trap p27 in the degradable state (see Figure 2I)^27, 28, 30^.

### Abl Requires an Arginine in the *α*D6 Residue, Abl^R328^, to Selectively Recognize pTyr in the -1 Position

We next sought to determine the biochemical basis for selectivity for pTyr priming in the -1 and +1 positions of the optimal substrates of Abl and Src. To identify candidate determinants of -1 pTyr recognition by Abl, *in* silico structural modelling was performed using a crystal structure of Abl in complex with its most optimal non-phosphorylated substrate Abltide (PDB: 2G2F, EAI-F-AAPF, where the F at position 4 is a non-phosphorylatable stand-in for tyrosine)^37^. The -1 isoleucine was substituted computationally with a pTyr, and a favorable rotamer manually selected. Hypothesizing that the negatively charged pTyr would be coordinated by positively charged amino acid residues in Abl, we searched for basic residues nearby. This identified Abl^R328^ as a promising candidate, particularly given its distance and relative position to a substrate - 1 pTyr and the fact that the guanidino group of Arg328 forms a potential electrostatic interaction (3.2 Å) with the phosphate group of the pTyr residue (Figures 3A and 3B). In Src, which does not show strong pTyr selection in the -1 position, and instead preferentially selects for isoleucine over pTyr, the corresponding amino acid at this position is a lysine. Mutation of R328 in Abl to lysine (Abl^R328K^) resulted in a kinase whose motif remained largely unchanged but demonstrated reduced selectivity for pTyr in the - 1 position relative to selection for isoleucine when assayed by PSPL (Figures 3C-D and S3A). Mutation of R328 to a non-basic alanine residue (Abl^R328A^) further decreased the selection for pTyr in the -1 position relative to Ile, while otherwise maintaining selectivity for all other aspects of the optimal substrate motif (Figures 3C-D and S3A). These results provide strong evidence that R328 (corresponding to the sixth amino-acid residue in alpha-helix D, and thus referred to as *α*D6), acts as a major determinant of recognition of N-terminal pTyr-primed substrates by the active site of Abl kinase.

**Figure 3.**
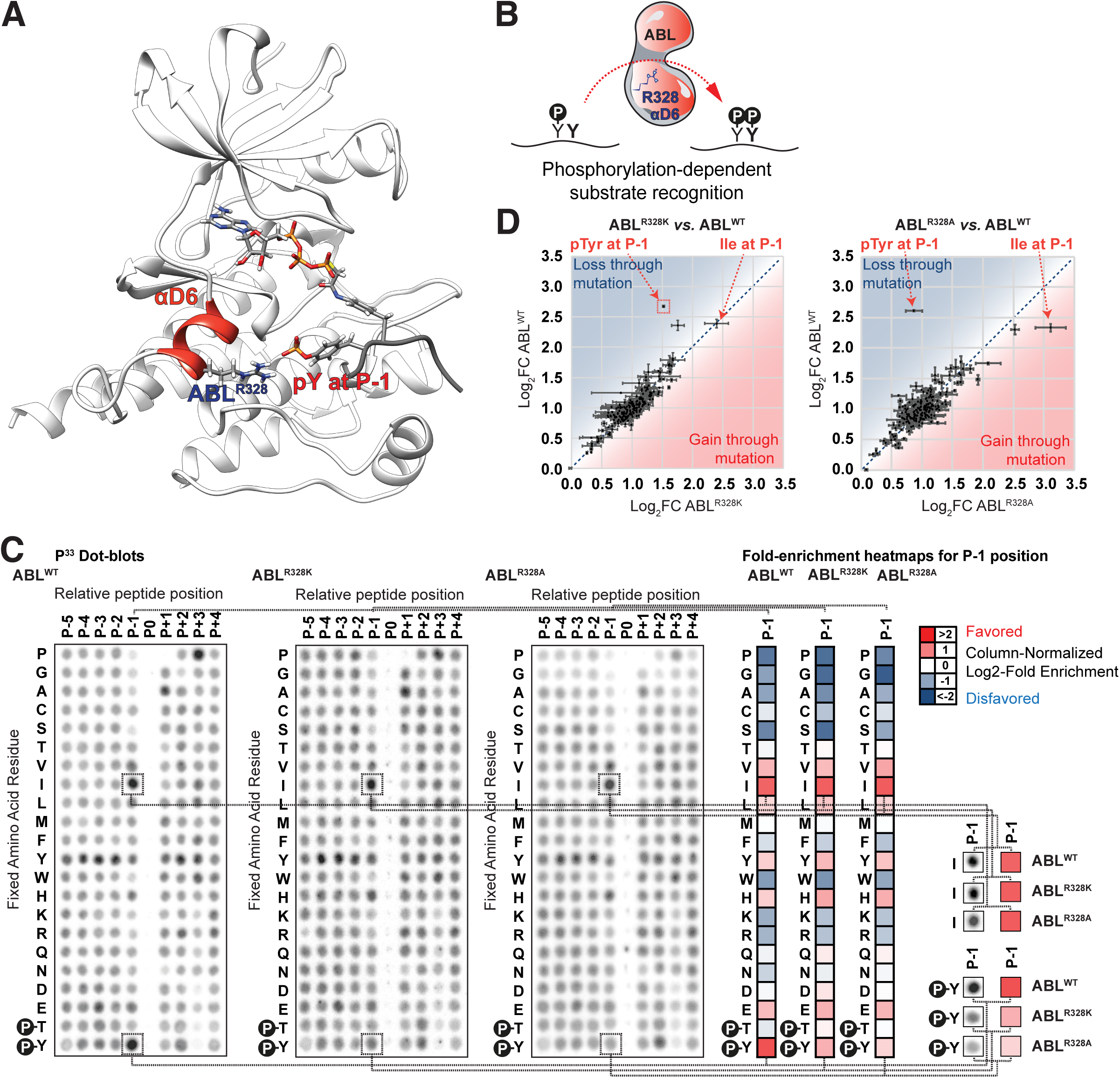
The Arginine in Position *α*D6, Abl^R328^, is Required for the N-terminal Recognition of pTyr in the Active Site of Abl Kinase. (A) Structural modeling of Abl and its optimal substrate peptide Abltide (PDB: 2G2F)^37^, where the canonical Ile directly N-terminal of the target tyrosine is replaced with pTyr, depicted in red, and the closest basic residue in Abl, Abl^R328^, is depicted in blue. (B) Graphical representation of the non-canonical, phosphorylation-dependent motif for Abl kinase with Abl^R328^ as a candidate molecular determinant driving the recognition of pTyr directly N-terminal from the target tyrosine. (C) Positional Scanning Peptide Library (PSPL) screening results for Abl^WT^, Abl^R328K^ and Abl^R328A^ kinases with the different fixed amino acid residues as rows (x axis) and different relative positions where the residues are fixed as columns (y axis). A column-normalized fold-enrichment heatmap for each kinase variant for the relative position directly N-terminal from the phosphorylation site position (P-1) is shown on the right with red denoting a favored amino-acid residue and blue denoting a disfavored one. (D) Scatterplots quantitatively comparing the column-normalized fold-enrichment values for each spot in the PSPL experiments comparing between Abl^R328K^ and Abl^WT^ (left), and Abl^R328A^ and Abl^WT^ (right). Datapoints in the upper-left region of the scatterplot indicate that a specific amino acid residue in a specific position has become less favored or more disfavored, whereas datapoints in the lower-right region of the scatterplot would indicate a change in specificity leading to a more favorable or less disfavorable effect for that amino acid in that position. See also Figure S3.

### Src Requires an Arginine in the *α*G2 Residue, Src^R472^, to Selectively Recognize pTyr in the +1 Position

A similar approach was used to identify the molecular determinants of pTyr recognition in the +1 position for Src. Since there are no entries in the PDB of Src kinase in complex with its optimal non-phosphorylated substrate peptide, we computationally superimposed the Src structure (PDB: 3D7T)^38^ to that of the Abl: Abltide complex (PDB: 2G2F)^37^ shown in Figure 3A. The +1 alanine of Abltide was substituted computationally with a pTyr, and a favorable rotamer selected. Examination of the resulting model revealed R472, as a promising candidate for making electrostatic interactions with the pTyr phosphate (Figures 4A-B). The corresponding amino acid in Abl, which does not show strong selection for pTyr selection in the -1 position, and instead preferentially selects for alanine, is a serine. Mutation of R472 in Src to serine (Src^R472S^) resulted in reduced selectivity for pY in the +1 position of the motif, although selectivity for the optimal non-phosphorylated residue in the +1 position, glycine, was also reduced.

**Figure 4.**
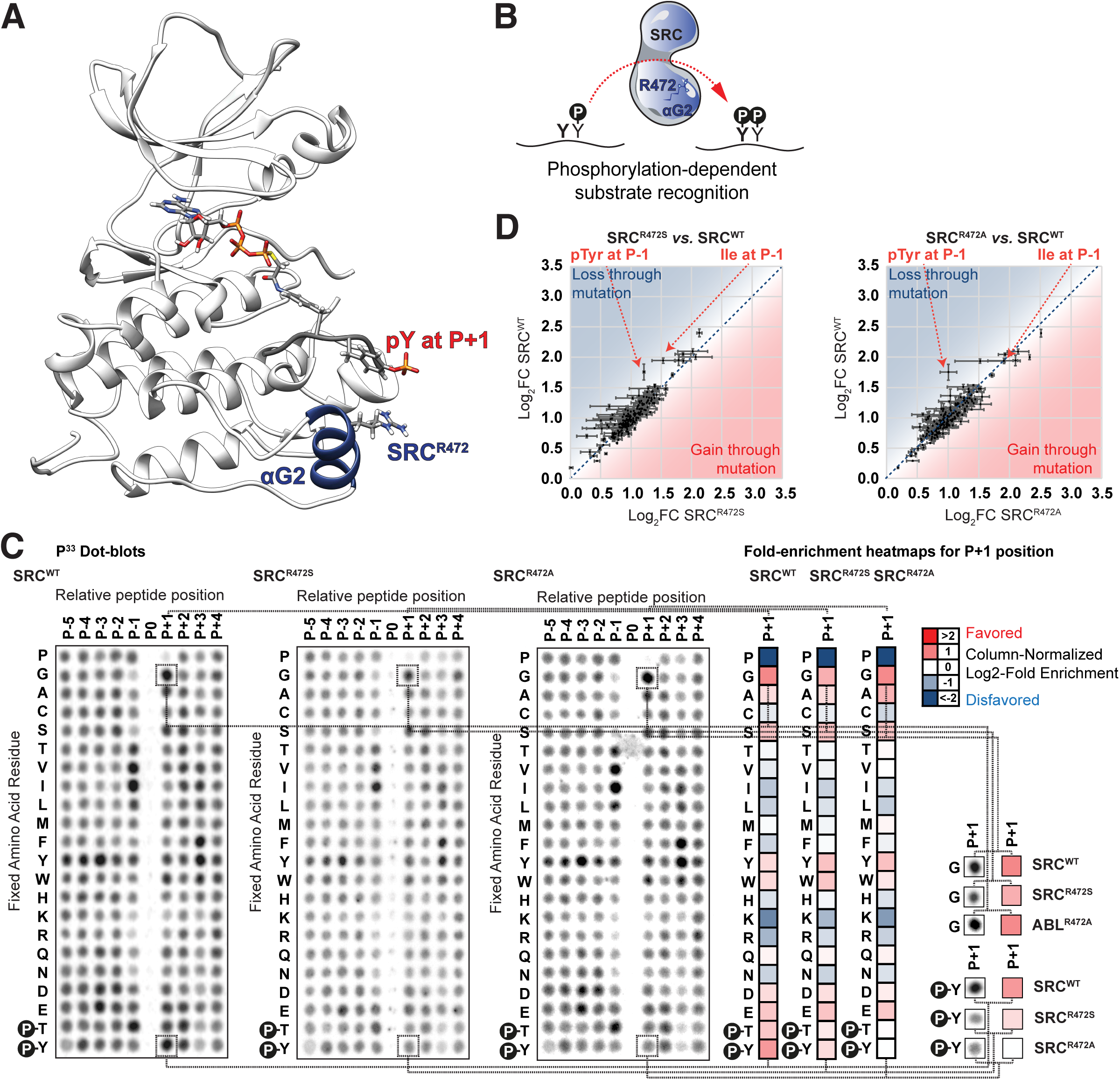
The Arginine in Position *α*G2, Src^R472^, is Required for the C-Terminal Recognition of pTyr in the Active Site of Src Kinase. (A) Structure of Src (from PDB: 3D7T)^38^ by homology modeling it to the structure of Abl with its optimal substrate peptide (from PDB: 2G2F)^37^, where the canonical residue directly C-terminal of the target tyrosine is replaced with pTyr, depicted in red, and the closest basic residue in Src, Src^R472^, is depicted in blue. (B) Graphical representation of the non-canonical, phosphorylation-dependent motif for Src kinase with Src^R472^ as a candidate molecular determinant driving the recognition of pTyr directly C-terminal from the target tyrosine. (C) Positional Scanning Peptide Library (PSPL) screening results for Src^WT^, Src^R472S^ and Abl^R472A^ kinases with the different fixed amino acid residues as rows (x axis) and different relative positions where the residues are fixed as columns (y axis). A column-normalized fold-enrichment heatmap for each kinase variant for the relative position directly C-terminal from the phosphorylation site position (P+1) is shown on the right with red denoting a favorable amino-acid residue and blue denoting a disfavorable one. (D) Scatterplots quantitatively comparing the column-normalized fold-enrichment values for each spot in the PSPL experiments comparing between Src^R472S^ and Src^WT^ (left), and Src^R472A^ and Src^WT^ (right). Datapoints in the upper-left region of the scatterplot indicate that a specific amino acid residue in a specific position has become less favored or more disfavored, whereas datapoints in the lower-right region of the scatterplot would indicate a change in specificity leading to a more favorable or less disfavorable effect for that amino acid in that position. See also Figure S4.

Mutation of R472 to alanine, however, completely eliminated +1 pTyr selection, with no effect on the strong selectivity for Gly (Figures 4C-D and S4A). Taken together, these results indicate that arginine 472, (corresponding to the second amino-acid residue in alpha-helix G, and thus referred to as *α*G2), acts as a major determinant of recognition of C-terminal pTyr-primed substrates by the active site of Src kinase.

### Multiple Tyrosine Kinases Use the *α*D6 and *α*G2 Molecular Determinants to Selectively Recognize -1 and +1 pTyr-primed Substrates

We extended our analysis of the two major determinants of pTyr priming, *α*D6 and *α*G2, to all other tyrosine kinases (Figures 5A, B and Figures S5A-C). This revealed that an arginine residue in the *α*G2 position, which drives C-terminal pTyr recognition in Src, is rarely present in the tyrosine kinome, and instead is largely restricted to Src and a few other closely related kinases. In contrast, an arginine residue in the *α*D6 position, which drives N-terminal pTyr recognition in Abl, is widely conserved in many other non-receptor and receptor tyrosine kinases. To test whether the pTyr recognition function is also conserved, we compared eleven tyrosine kinases by over-expressing and isolating them from HEK293T cells, followed by optimal substrate motif analysis using PSPL (Figure 5C). All of the kinases that contained a conserved arginine residue at their *α*D6 site, including the non-receptor tyrosine kinases Fes, Arg, Csk, and Brk and the receptor tyrosine kinases Alk and IRK, preferentially phosphorylated substrates with pTyr directly N-terminal from their target tyrosine. In contrast, all of the tyrosine kinases that lack an *α*D6 arginine residue, including the non-receptor tyrosine kinases Blk, Hck, and Lyn, showed no selectivity for a -1 pTyr in their optimal substrate motifs. Notably, these kinases also lack an arginine residue in their *α*G2 sites, and similarly, none of them displayed a preference for pTyr C-terminal to their target tyrosine, consistent with our structural motif determinant analysis (Figure 4).

**Figure 5.**
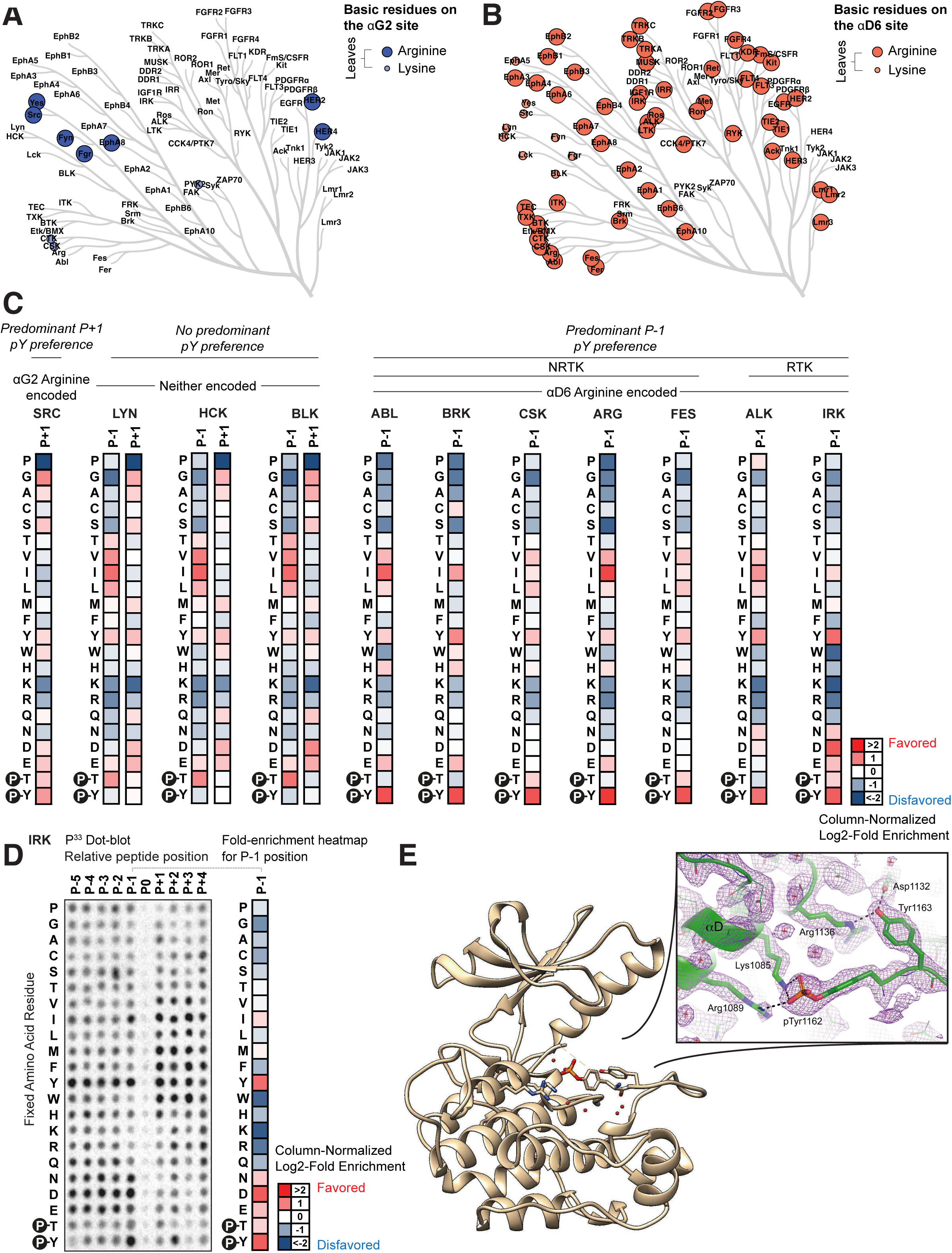
Multiple Tyrosine Kinases Recognise pTyr in Their Active Site and Encode the Same Molecular Determinants. (A) Dendrogram displaying the phylogenetic relationship of the human tyrosine kinases with leaves coloured according to the presence or absence of basic amino acid residues (arginine or lysine) at the *α*G2 sites. (B) Dendrogram displaying the phylogenetic relationship of the human tyrosine kinases with leaves coloured according to the presence or absence of basic amino acid residues (arginine or lysine) at the *α*D6 sites. (C) Column-normalized fold-enrichment heatmaps for different kinases quantifying the PSPL results for the relative position directly N-terminal (P-1) and/or C-terminal (P+1) from the phosphorylation site position, with red denoting a favored amino-acid residue and blue denoting a disfavored one. (D) Positional Scanning Peptide Library (PSPL) screening results for IRK with the different fixed amino acid residues as rows (x axis) and different relative positions where the residues are fixed as columns (y axis). A column-normalized fold-enrichment heatmap for the relative position directly N-terminal from the phosphorylation site position (P-1) is shown on the right with red denoting a favored amino-acid residue for IRK and blue denoting a disfavored one. (E) X-ray Crystal structure of bis-phosphorylated (pTyr1158, pTyr1162) IRK. Inset. The 2F_o_-F_c_ electron density (at 2.15 Å) is shown in purple, contoured at 1σ. Tyr1163 (P residue) of the activation loop is bound in the active site (in *cis*), hydrogen-bonded (black dotted lines) to Asp1132 and Arg1136 in the catalytic loop. pTyr1162 (P-1 residue) is salt-bridged (black dotted lines) to *α*D2 (Lys1085) and *α*D6 (Arg1089). Red crosses represent water molecules. See also Figure S5.

Interestingly, many tyrosine kinases, including the prototypical insulin receptor (IRK)^39^, contain consecutive tyrosine (YY) sites in their activation loops^40–44^. We wondered whether some of these kinases could undergo autophosphorylation and activation in a pTyr-primed manner. We focused on IRK (Figures 5C-E) because biochemical experiments have shown that, following insulin binding, the kinase domain is rapidly phosphorylated on Tyr-1162 and 1158, sequentially followed by slower autophosphorylation on Tyr-1163^43, 45, 46^. We purified and crystallized the kinase domain of IRK and solved the X-ray crystal structure of IRK at 2.15 Å resolution (Figure 5E).

Fortuitously, the kinase domain had undergone phosphorylation on the Tyr-1162 site prior to crystallization. Notably, in the resulting X-ray structure, direct interaction between this pTyr residue and the *α*D6 arginine residue (Arg1089) as well as an adjacent lysine residue (K1085), was observed.

### The Major Determinants of -1 and +1 pTyr Recognition Are Not Mutually Exclusive and Are Sufficient to Selectively Recognize Phosphoprimed Substrates

Having shown that an arginine residue in the *α*D6 or *α*G2 positions of Abl and Src, respectively are necessary for pTyr recognition in the -1 and +1 position, we next sought to determine whether arginine residues at those determinant sites are sufficient to acquire pTyr recognition function. To investigate this, we chose a tyrosine kinase, Frk, that contains non-basic residues at both its *α*D6 (Frk^A440^) and *α*G2 (Frk^Q319^) sites. Single and double arginine mutants (i.e., Frk^A440R^, Frk^Q319R^, and Frk ^A440R Q319R^) were generated, and the mutant recombinant proteins overexpressed, purified, and screened for the optimal phosphorylation motifs using PSPL (Figures 6A-B). Frk^A440R^ containing an arginine substitution for alanine in the *α*D6 position, resulted in marked pTyr selection in the -1 position of the phosphorylation motif. In contrast, Frk^Q319R^, containing an arginine residue in place of the natural glutamine in the *α*G2 position, led to a specific increase in the +1 recognition of pTyr compared to Frk^WT^. Surprisingly, a double mutant, Frk^A440R Q319R^, containing both the *α*D6 (Frk^A440R^) and *α*G2 (Frk^Q319R^) arginine sites, resulted in increased selection for pTyr at both the -1 and +1 positions of its optimal substrate motif relative to Frk^WT^ to approximately the same levels as that seen with the individual mutations alone (Figures 6B and S6A), indicating that these determinants are not mutually exclusive.

**Figure 6.**
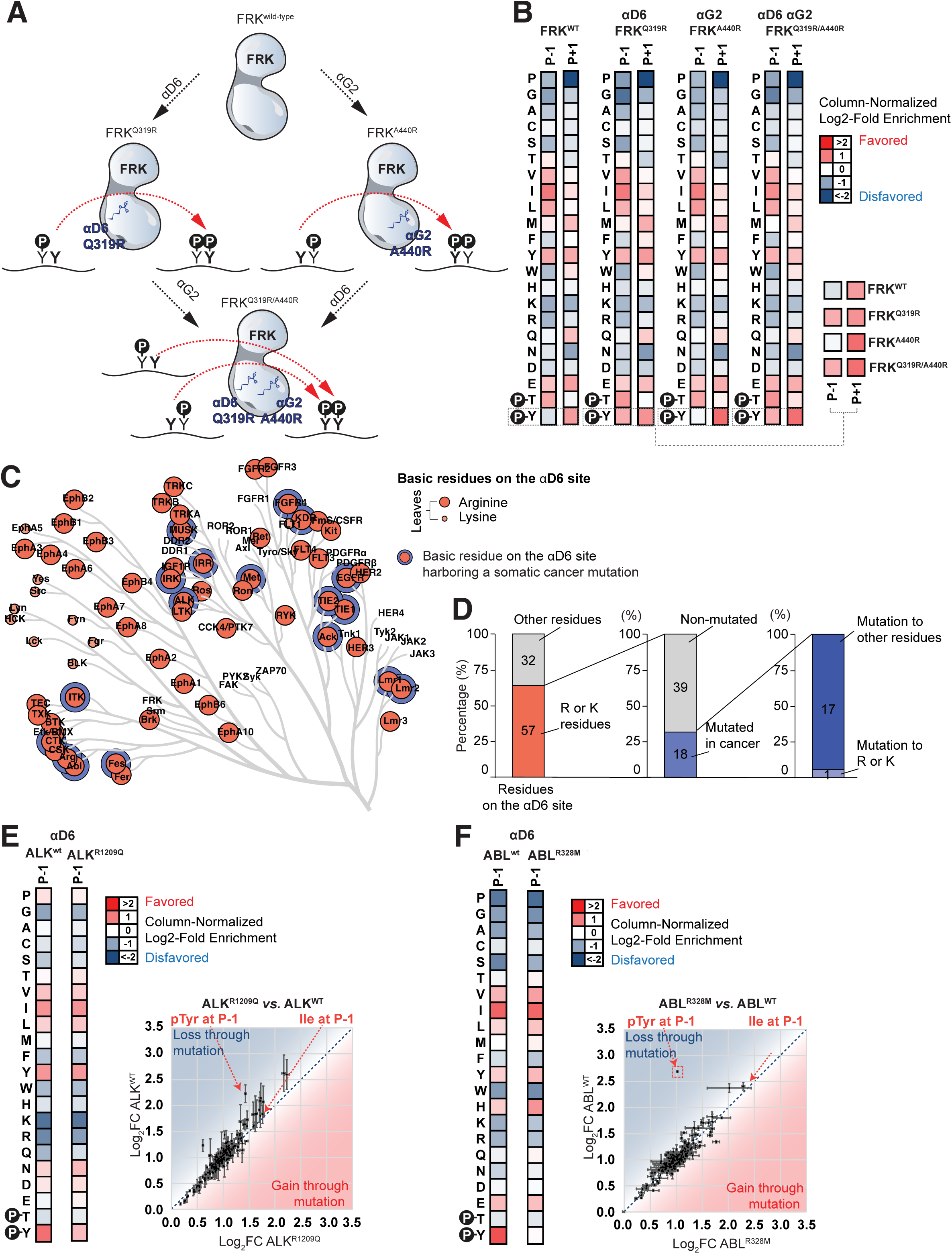
The Two Molecular Determinants are Non-Exclusive and Sufficient to Drive Active-Site Recognition of pTyr and Cancer Somatic Mutations Suppress this Recognition. (A) Graphical model illustrating our hypothesis that the addition of the basic residue arginine to the *α*D6 site (left), *α*G2 site (right) or both (bottom) would lead to pTyr recognition directly N-terminal, C-terminal or on both sides from the target tyrosine. (B) Column-normalized fold-enrichment heatmaps for the different FRK variants quantifying the PSPL results for the relative positions directly N-terminal (P-1) and C-terminal (P+1) from the phosphorylation site position, with red denoting a favored amino-acid residue and blue denoting a disfavored one. For easier comparison, the results for pTyr on these (P-1 and P+1) positions for the four FRK variants are re-plotted at the bottom right. (C) Dendrogram displaying the phylogenetic relationship of the human tyrosine kinases with its leaves coloured in red according to the presence of basic amino-acid residues (arginine, lysine or histidine) on the *α*D6 as well as in blue if a cancer somatic mutation has been observed on that same site. (D) Summary of residue composition and cancer mutation status at the αD6 site across tyrosine kinases. (E, F) Left. Column-normalized fold-enrichment heatmaps for the cancer somatic mutation Alk^R1209Q^ (E) or Abl^R328M^ (F) and its respective wild-type counterpart quantifying the PSPL results for the relative positions directly N-terminal (P-1) from the phosphorylation site position, with red denoting a favored amino-acid residue and blue denoting a disfavored one. Right. Scatterplot quantitatively comparing the column-normalized fold-enrichment values for each spot in the PSPL experiments comparing between Alk^R1209Q^ and Alk^WT^ (E) or Abl^R328M^ and Abl^WT^ (F). Datapoints in the upper-left region of the scatterplot indicate that a specific amino acid residue in a specific position has become less favored or more disfavored, whereas datapoints in the lower-right region of the scatterplot would indicate a change in specificity leading to a more favorable or less disfavorable effect for that amino acid in that position. See also Figure S6.

### Somatic Mutations in Cancer at the *α*D6 Site Abrogate Recognition of pTyr by Abl and Alk Kinases

As noted previously, 57 tyrosine kinases contain an arginine or lysine in the αD6 position (Figure 5A and Figures 6C, D). Of these, 18 have been found to contain cancer-associated non-synonymous mutations at the *α*D6 site, with 17 of these 18 mutations resulting in loss of a basic residue. We therefore selected one of these mutations in a non-receptor tyrosine kinase (Abl) and another one in a receptor tyrosine kinase (Alk) for further study. The Abl^R328M^ mutation was reported in a CML patient that was being treated with the Abl inhibitor Imatinib^47^. The Alk^R1209Q^ mutation has been reported more widely, including its presence in a brain metastasis from a patient with acute lymphoblastic leukemia^48^, as well as in cases of intestinal adenocarcinoma^49^, breast carcinoma^50^, lymph node metastases from malignant melanoma^51^, and non-small cell lung cancer following treatment with the Alk inhibitor Alectinib^52^. The Alk^R1209Q^ and Abl^R328M^ mutants were expressed, purified, and screened for motif selectivity using PSPL. The R1209Q mutation in Alk substantially reduced selection for pTyr in the -1 position (Figure 6E and Figure S6B), while the R328M mutation in Abl completely eliminated it (Figure 6F and Figure S6C).

### Non-canonical pTyr Encoding Substrates Downstream of Bcr-Abl Drive Anti-Oncogenic Molecular Functions *In Vivo*

To further explore the effects of alanine and methionine mutations involving the αD6 residue in Abl (Figures 3C, D, and Figure 6F) on canonical and non-canonical motif preferences, particularly in the setting of one its natural substrates, CrkII^53^, we performed kinase assays using four different peptide variants encompassing the sequence around Tyr 221. In these variants the Pro residue in the -1 position of wild-type CrkII was replaced by Ile, pTyr, or Phe (Figure 7A and Figure S7A). Mutation of Pro 220 to either pTyr or Ile, but not Phe, resulted in enhanced levels of peptide phosphorylation using wild-type Abl. Replacement of Abl’s αD6 arginine with either alanine or methionine substantially reduced the phosphorylation of the pTyr CrkII peptide variant, while having no significant effect on the isoleucine variant. This further confirmed the importance for phosphopriming selection of a basic residue in this position in Abl substrates, but not for canonical motif selection.

**Figure 7.**
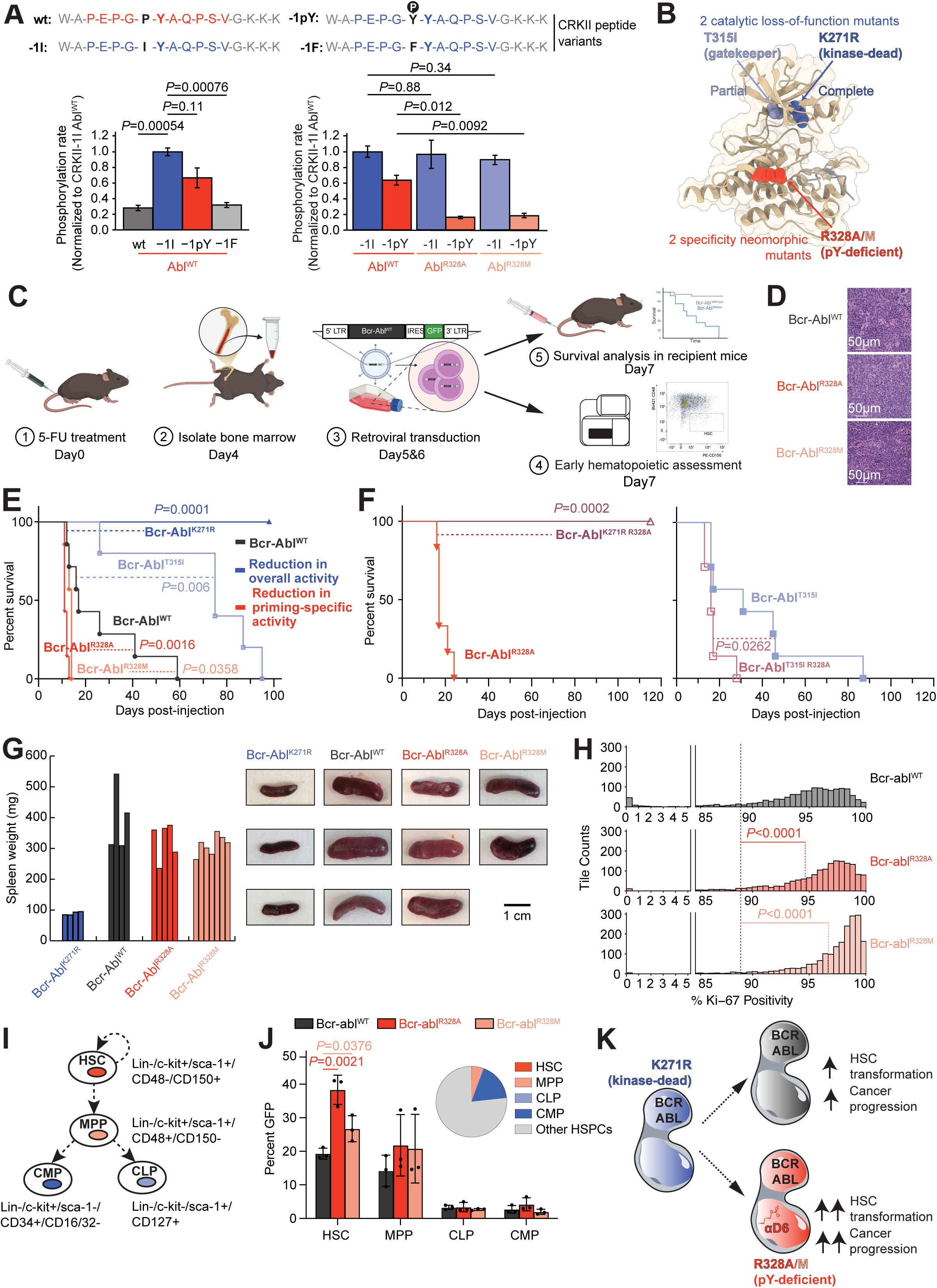
The Bcr-Abl^R328A^ and Bcr-Abl^R328M^ Variants Result in High Stem Cell Progenitors and Faster Myeloproliferative Progression Uncovering an Anti-Oncogenic Function Downstream of the Pro-Oncogenic Bcr-Abl Kinase *In Vivo*. (A) Phosphorylation rates of CrkII peptide variants with substitutions at the −1 position relative to the target tyrosine (illustrated at the top). Left. Phosphorylation rates measured for Abl^WT^ using −1F, −1I, and −1pY variants, normalized to −1I. Right. Relative phosphorylation rates comparing the kinase activity of Abl^WT^, Abl^R328A^ and Abl^R328M^ against two variant peptides of the Abl natural substrate CrkII. Data represent reaction slopes from in vitro kinase assays shown in Figure S7A. (B) Structural illustration of the 2 catalytic loss-of-function mutants, Bcr-Abl^T315I^ and Bcr-Abl^K271R^, and 2 specificity neomorphic mutants, Bcr-Abl^R328A^ and Bcr-Abl^R328M^ incapable of recognizing phosphoprimed substrates. (C) Schematic representation of Bcr-Abl-driven *in vivo* mouse model experiments. (D) Sections from spleen of representative Bcr-Abl^WT^, Bcr-Abl^R328A^, and the Bcr-Abl^R328M^ transplanted animals were stained with haematoxylin and eosin (H&E). Scale bars are 50 μm. (E) Comparison of the percent survival over time for mice harbouring Bcr-Abl^WT^, Bcr-Abl^T315I^, Bcr-Abl^K271R^, Bcr-Abl^R328A^, and the Bcr-Abl^R328M^ variants. (F) Left. Comparison of the percent survival over time for mice harbouring the hypomorphic gatekeeper variant Bcr-Abl^T315I^ either by itself or in combination with the Bcr-Abl^R328A^ mutation, Bcr-Abl^R328A T315I^. Right. Comparison of the percent survival over time for mice harbouring the variant Bcr-Abl^R328A^ either by itself or in combination with the kinase-dead mutation, Bcr-Abl^R328A K271R^. (G) Left. Spleen weights measured from mice injected with Bcr-Abl^K271R^, Bcr-Abl^WT^, Bcr-Abl^R328A^, or Bcr-Abl^R328M^. Right. Representative images of spleens from mice injected with Bcr-Abl^K271R^, Bcr-Abl^WT^, Bcr-Abl^R328A^, or Bcr-Abl^R328M^. (H) Tile-based quantification of Ki-67^+^ cells in spleen IHC sections from mice expressing Bcr-Abl^WT^, Bcr-Abl^R328A^, or Bcr-Abl^R328M^. Ranges 0–5% and 85–100% are shown; the full distribution is provided in Figure S7D. (I) Diagram illustrating cell surface markers and relationship among the different stem cells and hematopoietic progenitors in terms of self-renewal (curved arrow) and developmental fate decisions (straight arrows) during differentiation. HSC, Hematopoietic Stem Cells; MPP, Multi-Potent Progenitor; CLP, Common Lymphoid Progenitor; CMP, Common Myeloid Progenitor. (J) Left. Comparison of the percentage of GFP-high cells within progenitor subpopulations including HSCs, MPPs, CLPs, and CMPs, following transduction of total HSPCs with Bcr-Abl^WT^, Bcr-Abl^R328A^, or Bcr-Abl^R328M^. Data are shown as bar plots with individual data points and medians (n = 3 mice per group). Right. The pie chart shows the distribution of these subpopulations within the Bcr-Abl^WT^-transduced HSPC pool. Equivalent distributions for Bcr-Abl^R328A^ and Bcr-Abl^R328M^ are shown in Figure S7E. (K) Graphical model summarizing our findings, where variants of Bcr-Abl impaired in their ability to recognize pTyr directly N-terminal from their target tyrosine site (Bcr-Abl^R328A/M^) lead to a higher percentage of transformed, Bcr-Abl-GFP-high hematopoietic stem cells and faster myeloproliferative progression *in vivo* than WT Bcr-Abl. Our results imply that non-canonical pTyr-driven substrates downstream of Bcr-Abl^WT^ suppress transformation of hematopoietic stem cells and myeloproliferative progression *in vivo*. See also Figure S7.

Since the principal consequence of these Abl mutations is a loss of selectivity for phosphorylating -1 pTyr-primed substrates (Figures 3C, D, Figure 6F, Figure 7A and Figure S7A), we next studied these mutations in the context of the highly cancer-relevant wild-type Bcr-Abl fusion product, alongside Bcr-Abl fusions involving a kinase-dead Abl, and a hypomorphic kinase-mutation in Abl at the gatekeeper residue, T315I^54^ (Figure 7B and Figure S7B), using a mouse model of chronic myeloid leukemia (CML)^55^.

In this model (Figure 7C), hematopoietic stem and progenitor cells (HSPCs) are extracted from the bone marrow of donor mice, expanded *in vitro*, and retrovirally transduced twice, 24 h apart, with wild-type or mutant Bcr-Abl constructs^55^. Lethally irradiated recipient mice underwent bone marrow reconstitution with these HSCs 24h after final transduction, using seven mice per construct and an equal number of transduced cells in each animal.

Leukemia developed in all mice except those receiving the kinase-dead Bcr-Abl fusion product (Figure 7D). The aggressiveness of the disease appeared to directly scale with kinase activity – mice receiving wild-type Bcr-Abl all succumbed within 60 days, while those receiving the gatekeeper mutant hypomorph survived up to about 95 days. We expected that mice receiving *α*D6-mutant phosphopriming-deficient neomorphic Bcr-Abl constructs, maintaining most but not all of the biochemical substrate specificity of wild-type Bcr-Abl (Figure 3D and Figure 6F), would show intermediate survival between the wild-type and the T315I hypomorph. Surprisingly, however, these mice instead died more quickly than those receiving wild-type Bcr-Abl (Figure 7E, log-rank test: *P* = 0.0016 for R328A; *P* = 0.0358 for R328M), dying in 15 or fewer days. To verify that the accelerated leukemic killing effect of these neomorphic mutations depended on intrinsic Abl kinase activity, we generated compound mutant in which the *α*D6 R328A mutation was combined with the K271R kinase-dead mutation (Figure 7F, left panel).

None of these mice developed leukemia, verifying that the neomorphic killing effect requires Abl kinase activity. Furthermore, addition of the neomorphic R328A mutation to the T315I hypomorphic mutation accelerated leukemogenic death in these animals, relative to the T315I mutation alone (Figure 7F, right panel), confirming that the effect of these *α*D6 mutations is mediated by changes in the motif specificity of the Abl kinase domain.

Leukemic infiltration of the spleen, as assessed by spleen weight, was similar at the time of sacrifice in each cohort, despite the shorter survival time in the cohort bearing phosphopriming mutant Bcr-Abl, suggesting an increased rate of leukemic cell proliferation in these animals (Figure 7G). Consistent with this, splenic sections from these animals demonstrated a higher proportion of Ki-67^+^ cells, compared to animals bearing wild-type Bcr-Abl (Figure 7H and Figures S7 C-D).

To further explore this, we characterized the transduced bone marrow cell populations 24h after final transduction (Figure 7C). The proportion of Bcr-Abl fusion expressing cells in each hematopoietic progenitor subtype was quantified by GFP positivity. A significantly enhanced percentage of GFP-positive cells was specifically observed for the phosphopriming mutant Bcr-Abl fusion proteins relative to wild-type Bcr-Abl in hematopoietic stem cells (HSCs). As these are the well-established cells of origin and propagation for CML, this suggests a survival advantage. No significant difference was observed in the percentage of GFP positive cells in the multipotent progenitor (MPP), common myeloid progenitor (CMP), or common lymphocytic progenitor (CLP) populations (Figures 7I-J and Figure S7 E). Altogether, our results support a model where specific defects in recognizing pTyr by Bcr-Abl lead to increased percentages of transformed HSC, and a more aggressive disease (Figure 7K).

## Discussion

In this manuscript, we investigated the molecular determinants of tyrosine phosphopriming for tyrosine kinase substrate selection by tyrosine kinases, and its implications for biological signaling networks. The concept that a kinase might require prior phosphorylation at an adjacent residue in order to phosphorylate a cognate substrate has best been demonstrated for the serine/threonine kinase GSK3, which has been shown to phosphorylate substrates with a phosphorylated serine or threonine residue 4 positions C-terminal to the GSK3 substrate phosphorylation site^56^. In addition, we and others have discovered that the tyrosine kinases EGFR and BMX can utilize phosphopriming on adjacent tyrosine residues as a determinant of substrate specificity^17, 19^. Moreover, we have recently demonstrated that up to 60% of the tyrosine kinome favors a phosphoserine (pSer), phosphothreonine (pThr) or pTyr within the -5 to +4 positions relative to their substrate tyrosine^20^, suggesting a much more common role for this relatively novel form of inter-kinase interaction in the construction of biological signaling networks.

Protein kinase signaling networks are frequently depicted as using writers (kinases), readers (Phosphopeptide-Binding Domains (PBDs)), and erasers (phosphatases) as a means to communicate information^6^. Some serine/threonine kinases embody two of these roles, using PBDs such as forkhead-associated (FHA) domains and the Polo-box domain of the Polo-like kinases to assist in targeting the kinase to its substrates, which have been previously phosphorylated at a relatively distal site^57, 58^. Similarly, in the case of non-receptor tyrosine kinases, pTyr recognition by an adjacent SH2 domain appears to be particularly prevalent, since ∼78% of these contain an SH2 domain^59^. Our findings that many tyrosine kinase catalytic domains themselves recognize pTyrs within their optimal substrate phosphorylation motif, without the need for a distinct phosphopeptide binding domain, blends these otherwise binary definitions of reader and writer modules.

Src and Abl are examples of non-receptor tyrosine kinases that can utilize this PBD-independent-mechanism of pTyr recognition to create a signaling pathway topology of potential mutual interdependency, since the action of each kinase alone on a tandem YY motif would create an optimal substrate for the other or even itself. We show that this mutuality may converge on the substrate p27, which is phosphorylated at residues Y88 and Y89 to control G1 arrest, possibly through kinetic trapping (Figure 2I). This mechanism, or another that relies on phosphopriming and sequential phosphorylation, may help clarify previous observations implicating Y88, Y89, Abl and Src in p27 control^14, 16, 28–30^.

Using Abl and Src as model kinases, we identified the *α*D6 and *α*G2 positions within the kinase domain as critical determinants of phosphopriming recognition at the -1 and +1 substrate sites, respectively. These determinants, when substituted into the kinase domain of Frk, were sufficient to enhance specificity for phosphoprimed substrates in the -1 or +1 positions. Intriguingly, 64% of tyrosine kinases contain a lysine or arginine residue at the *α*D6 position, while 11% have a lysine or arginine in *α*G2, indicating that phosphopriming may be broadly utilized within the tyrosine kinome, generally in good agreement with the results of a comprehensive screening for substrate specificity of all tyrosine kinases^20^.

In 17 of the 57 kinases bearing the *α*D6 determinant of -1 tyrosine phosphopriming, mutation of this *α*D6 basic residue to one less compatible with -1 tyrosine recognition has been observed. While the contribution of these mutations to the cancer phenotype is unclear, we show here in the case of Bcr-Abl that a mutation disrupting phosphopriming recognition resulted in a more aggressive development of leukemia *in vivo* (Figure 7). These surprising results suggest that, in contrast to the current dogma in oncology, where substrates downstream of an oncogenic kinase are typically considered to primarily drive pro-oncogenic functions, non-canonical substrates containing pTyr-primed sequences may alternately drive anti-oncogenic functions (Figure 7K).

Tyrosine kinase signaling networks have traditionally been interpreted in light of single site phosphorylation and single phosphosite recognition. By contrast, the breadth of phosphopriming recognition potentially available within the tyrosine kinome, as shown here and in our previous works^17, 19, 20^, highlights the possibility of a much more nuanced and dynamic regulation of signaling networks through dual pTyr motifs, including but not limited to tandem YY sequences. Such multisite tyrosine phosphorylation could be used, for example, to generate dynamically controlled “AND”, “OR”, or “NOR” gates, in which a specific biological outcome depends on the activity or inactivity of one kinase acting before or after another to post-translationally modify a shared kinase substrate sequence. Indeed, our finding of a convergence between Src and Abl to regulate p27 levels is consistent with such a model, as is the discovery that certain individual SH2 domains, such as those in Vav, PLCg1, and Src itself can specifically recognize short peptide sequences containing two closely spaced pTyr residues with high affinity^60–64^.

The general extent to which these types of dual tyrosine phospho-modification and recognition are used to control complex cellular decisions should become more apparent in the future, particularly as emerging top-down phosphoproteomics mass spectrometry techniques are used to characterize signaling pathways in greater molecular detail^65^.

## Author contributions

P.C. and M.B.Y. conceived the study. P.C., N.C., L.R.M.B., M.C., A.R.B., Y.W.K., J.L., C.K.L., J.C., C.L., G.S., T.J.G.R., B.K., J.L.J. and S.H. designed and performed *in vitro*, structural and/or *in vivo* experiments, with supervision from P.C., C.B., L.C., R.C., J.P., B.H., M.H., S.H. B.A.J., and M.B.Y. S.S., and E.M.S., performed and analysed Mass Spectrometry-based experiments. P.C., and M.C. structurally modelled candidate determinants of substrate specificity *in silico*. P.C., B.A.J., and M.B.Y. wrote the manuscript with input from the other authors.

## Acknowledgements

We are grateful to Ian Cannell, Anthony Coyne, Simona Valeviciute, Daniel Lim, Robert Grant, Alex Kruswick, Karl Merrick along other current and past members of the Creixell and Yaffe laboratories, members of the MIT Center for Precision Cancer Medicine, Forest White, Ishwar Kohale, Benjamin E. Turk, Chad J. Miller, Rune Linding, Craig Simpson, Benedict-Tilman Berger, Stefan Knapp, Billie G. C. Griffith and Margaret C. Frame for their help, sharing reagents, discussions and advice leading up to this manuscript. This research was supported by a BBSRC Strategic LoLa grant (BB/MM00354X/1) (A.R.B. and C.B.); by the Jane Coffin Childs Memorial Fund, the Research Council of Finland, and the Finnish Cancer Foundation (J.L.); by the Claudia Adams Barr Program for Cancer Research award (J.L.J.); by an HHMI Gilliam Fellowship GT15758 (T.J.G.R.); by National Institutes of Health grants R35-CA197588, P01-CA120964, and P01-CA117969 (L.C.C.); by NIH/NCI R01 CA196703-01 (R.C.); by National Institute of Health grants R01-GM104047, R01-ES015339 and R35-ES028374 (M.B.Y.); by the Charles and Marjorie Holloway Foundation (M.B.Y.); and by the MIT Center for Precision Cancer Medicine; by the National Cancer Institute of the National Institutes of Health under Award Number K99CA226396 (P.C.), core support from Cancer Research UK (CRUK C9545/A29580) (P.C.) and the Children’s Brain Tumour Centre of Excellence (C9685/A26398) (P.C.). Support was also provided by the Cancer Center Support Grant P30-CA14051 from the National Cancer Institute and the Center for Environmental Health Sciences Support Grant P30-ES002109 from the National Institute of Environmental Health Sciences.

## Methods

### Molecular cloning

Most wild-type protein tyrosine kinases were kind gifts from Karen Colwill, Tony Pawson, Craig Simpson, and Rune Linding cloned into the Gateway destination vector including a C-terminal 3xFLAG epitope tag, V1900, by Gateway cloning (ThermoFisher). We made all point mutants by site-directed mutagenesis using Q5 Site-Directed Mutagenesis Kit (New England Biolab, NEB) and following its standard protocol.

### Protein over-expression, and isolation

All kinases used for PSPL screening were purified from transiently transfected HEK 293T cells, modifying protocols described in van de Kooij et al^66^ as required for tyrosine kinases. Briefly, 7 × 10^6^ HEK 293T cells were plated in 15cm plates, and the next day transfected with 20–25 µg kinase-DNA using polyethyleneimine (PEI). After overnight incubation, we replaced the transfection medium with fresh medium. 24 hr later, we washed all cells with cold PBS and harvested them by scraping in lysis buffer (20 mM Tris-HCl pH 7.5, 150 mM NaCl, 1 mM EDTA, 1% Triton-X100, 1 mM DTT, and 0.1 mM NaVO_3_) supplemented with protease inhibitors (complete EDTA-free protease inhibitor cocktail, Roche) and phosphatase inhibitors (PhosSTOP, Roche). Lysates were incubated on ice for 20 min, and cleared by centrifugation. Next, anti-FLAG M2-affinity gel (Sigma) was added to the cleared lysate, followed by incubation for 2 hr at 4°C while rotating. After incubation, the beads were pelleted by centrifugation, washed twice with lysis buffer and washed twice again with wash buffer (50 mM HEPES pH 7.4, 100 mM NaCl, 1 mM DTT, 0.01% NP-40, 10% glycerol). Protein was eluted from the beads by incubation for 1 hr at RT (while rotating) with 250 µl wash buffer supplemented with phosphatase inhibitors (PhosSTOP, Roche) and 0.5 mg/mL 3xFLAG peptide (APExBIO). To determine concentration and purity, a BSA standard curve and a sample of the eluate were analysed by SDS-PAGE and Coomassie-staining (SimplyBlue Safe stain, Life Technologies). Eluted kinase was snap-frozen and stored at −80°C.

### Positional Scanning Peptide Library (PSPL) Screening

Oriented Peptide Library Screens were performed as described in van de Kooij *et al*^66^, which followed a similar approach as Hutti *et al*^67^. but with a slightly adapted protocol so that the assay could be done in 1536-well plates, except with modifications as required for tyrosine kinases. The tyrosine-peptide library (custom order, JPT Peptide Technologies GmbH) consisted of 198 peptide pools with the sequence G-A-Z-X-X-X-X-Y-X-X-X-X-A-G-K-K-(LC-Biotin)-NH_2_, were X = mixture of all natural amino acids except Cys, Ser, Thr and Tyr, and LC-Biotin = long chain biotin. In each peptide pool, the Z is fixed to be a single amino acid, which can be any of the 20 natural amino acids or pThr or pTyr. In the example, the Z is positioned on the P-5 relative to central Y, but in the complete library the Z would be fixed in a position ranging from −5 to +4 relative to the central Y. Hence, there are nine different positions and 22 different amino acids, generating 198 different peptide pools. All peptides were dissolved in DMSO to 5 mM, arrayed in a 1536-well plate by multi-channel pipetting, and diluted to 500 µM with water.

For the kinase assay, 2.2 µl kinase buffer per well was dispensed in 1536-well plates using a Mantis nanodispenser robot (Formulatrix). The core kinase buffer consisted of 50 mM Tris-HCl pH 7.5, 0.1 mM EGTA, 1 mM DTT, 50 µM ATP, 1 mM NaVO_3_, 5 mM BGP and 0.1% Tween-20, and was supplemented with 10 mM MgCl_2_ and 2 mM MnCl_2_. Next, 250 nl peptide was transferred from the peptide-stock plate to the kinase assay plate using a pintool with slot tips (V and P Scientific, Pin type FP3S200). Subsequently, 200 nl of a mix of kinase and [γ-^33^P)-ATP was added using the Mantis, with a final concentration in each well ranging from 5 to 50 nM of the kinase and 0.027 μCi/μl for the [γ-^33^P]-ATP. Between every step, the 1536-well plate was centrifuged briefly to collect all liquid, and cooled on ice as needed to prevent evaporation. After addition of the kinase, the plate was incubated at 30°C for 2 hr. Subsequently, 250 nl of each reaction was spotted on streptavidin-membrane (Promega) using the pintool, the membrane was washed and dried, exposed to a phosphorimaging screen, and imaged on a Typhoon 9400 imager (GE Healthcare).

Spot intensities were quantified using ImageStudio (Li-COR BioSciences), and normalized by dividing the measured intensity for an individual spot by the average spot intensity for all amino acids in that position. For each kinase, OPLS-experiments were done at least twice, and normalized values were averaged for all replicates. Next, normalized values were log2-transformed in Microsoft Excel.

### *In vitro* kinase activity assays and determination of kinetic parameters

*In vitro* kinase assays with a peptide-substrate were performed as described in van de Kooij *et al*^66^. 5–50 nM of kinase was pre-incubated at 30°C for 30 min in the same kinase buffer as used for OPLS, but without the 0.1% Tween-20. Subsequently, [γ-^33^P]-ATP was added to a final concentration of 0.075–1 μCi/μl, and peptide added to a final concentration of 50 μM. Peptides were synthesized in-house at the Swanson Biotechnology Center – Biopolymers and Proteomics Facility at the 5 μmol starting resin scale per 96-well plate well, using an MultiPep small scale peptide synthesizer (Intavis) and standard FMOC chemistry. In addition to the peptide-specific sequences indicated, each peptide contains an N-terminal W-A sequence to allow concentration determination of peptide-solutions by UV-spectrophotometry, and a C-terminal G-K-K-K sequence to increase solubility and interaction with phosphocellulose. After adding substrate and [γ-^33^P]-ATP, the mixture was incubated at 30°C for 20 min, and every 5 min a sample was spotted on p81 phosphocellulose papers (Reaction Biology Corp.).

After spotting, the phosphocellulose papers were immediately soaked in 0.5% phosphoric acid. At the end of the assay, all papers were washed three times in 0.5% phosphoric acid, air-dried, transferred to vials with scintillation counter fluid and counted by a LS 6500 scintillation counter (Beckman Coulter). The resulting data points were analysed by linear regression, and the phosphorylation rate was defined as the slope of the linear regression curve.

### Targeted Mass-Spectrometry

#### Sample Preparation

Lysate preparation and digestion was done according to Kulak *et al*^68^. Briefly, cells were lysed using 20μl of lysis buffer (consisting of 6 M Guanidinium Hydrochloride, 10 mM TCEP, 40 mM CAA, 50 mM HEPES pH8.5). Samples were boiled at 95°C for 5 minutes, after which they were sonicated on high for 3x 10 seconds in a Bioruptor sonication water bath (Diagenode) at 4°C. After determining protein concentration with Bradford (Sigma), 50μg was taken forward for digestion. Samples were diluted 1:3 with 10% Acetonitrile, 50 mM HEPES pH 8.5, LysC (MS grade, Wako) was added in a 1:50 (enzyme to protein) ratio, and samples were incubated at 37°C for 4hrs. Samples were further diluted to 1:10 with 10% Acetonitrile, 50 mM HEPES pH 8.5, trypsin (MS grade, Promega) was added in a 1:100 (enzyme to protein) ratio and samples were incubated overnight at 37°C. Enzyme activity was quenched by adding 2% trifluoroacetic acid (TFA) to a final concentration of 1%. Prior to mass spectrometry analysis, the peptides were desalted on in-house packed C18 Stagetips^69^. For each sample, 2 discs of C18 material (3M Empore) were packed in a 200μl tip, and the C18 material activated with 40μl of 100% Methanol (HPLC grade, Sigma), then 40μl of 80% Acetonitrile, 0.1% formic acid. The tips were subsequently equilibrated 2x with 40μl of 1%TFA, 3% Acetonitrile, after which the samples were loaded using centrifugation at 4,000x rpm. After washing the tips twice with 100μl of 0.1% formic acid, the peptides were eluted into clean 500μl Eppendorf tubes using 40% Acetonitrile, 0.1% formic acid. The eluted peptides were concentrated in an Eppendorf Speedvac, and re-constituted in 1% TFA, 2% Acetonitrile prior to spike-in with synthetic, heavy-labeled peptides for Mass Spectrometry (MS) analysis.

#### MS data acquisition

For each sample, peptides were loaded onto a 2cm C18 trap column (ThermoFisher 164705), connected in-line to a 50cm C18 reverse-phase analytical column (Thermo EasySpray ES803) using 100% Buffer A (0.1% Formic acid in water) at 750bar, using the Thermo EasyLC 1000 HPLC system, and the column oven operating at 45°C. Peptides were eluted over a 70 minute gradient ranging from 6 to 60% of 80% acetonitrile, 0.1% formic acid at 250 nl/min, and the Q-Exactive HF-X instrument (Thermo Fisher Scientific) was run in targeted Parallel Reaction Monitoring (PRM) mode, focusing on the doubly phosphorylated peptide as heavy labelled standard and as unlabeled endogenous peptide. Full MS spectra were collected at a resolution of 120,000, with an AGC target of 3×10^6^ or maximum injection time of 50ms and a scan range of 350–1750 m/z. The MS2 spectra were obtained at a resolution of 30,000, with an AGC target value of 2×10^5^ or maximum injection time of 500ms, and normalised collision energy of 30. MS performance was verified for consistency by running complex cell lysate quality control standards, and chromatography was monitored to check for reproducibility. PRM data analysis was conducted in Skyline^70^, and the heavy standard reference was used for localizing the endogenous peptide on the gradient.

### p27 experiments

#### hTert-RPE1 Cell Culture

hTert-RPE1 mRuby-PCNA cells were from Joerg Mansfeld (ICR, London)^33^ and were maintained in DMEM (Gibco) + 10% FBS (Sigma) and 1% Penicillin-Streptomycin (P/S; Gibco) at 37°C and 5% CO2. Cell lines were checked by PCR on a monthly basis for absence of mycoplasma.

#### Generating p27-GFP tagged hTert-RPE1 mRuby-PCNA cell lines

The endogenous *CDKN1B* gene was tagged at the C-terminus using CRISPR-mediated gene tagging. The gRNA 5’-TCAAACGTAAACAGCTCGGT<u>GGG</u>-3’ (PAM site underlined) covering the *CDKN1B* stop codon was selected using crispr.mit.edu. Forward and reverse oligos for the gRNA were annealed and ligated into the BbsI cut pX330 bicistronic Cas9 and sgRNA expression plasmid pX330-U6-Chimeric_BB-CBh-hSpCas9 was a gift from Feng Zhang (Addgene plasmid # 42230)^71^.

For the homology donor plasmid, we PCR amplified *CDKN1B* homology arms from hTert-RPE1 genomic DNA using the high-fidelity Q5 DNA polymerase (New England Biolab, NEB). To PCR amplify the left homology arm (LHA) we used the forward primer with a NotI site: 5’-gcggccgcTAGAGGGCAAGTACGAGT-3’ and the reverse primer with a SalI site: 5’-gtcgacCGTTTGACGTCTTCTGAG-3’. To PCR amplify the right homology arm (RHA) we used the forward primer with a SpeI site 5’-actagtACAGCTCGgtgg<u>C</u>ttgatc-3’ (underlined base represents mutated base to change the PAM site in the recombined gene) and the reverse primer with a NotI site: 5’-gcggccgcctggggactgaattattca-3’. eGFP cDNA was PCR amplified from peGFP-C1 vector (Clontech) with the forward primer with a SalI site and a poly-glycine linker: 5’-gtcgacggaggaggagtgagcaagggcgaggag-3’ and the reverse primer with a SpeI site: 5’-actagtttacttgtacagctcgtc-3’. LHA, RHA and eGFP PCR products were ligated into the blunt-end cloning vector pJet1.2 (Fermentas) for restriction digest. LHA, RHA and eGFP digested inserts were ligated into NotI cut pAAV-MCS vector by 4-way ligation at a ratio of vector:inserts of 1:2:2:2, using T4 DNA ligase (NEB). All constructs were checked by Sanger sequencing before transfection into cells.

To generate Y88F and Y89F mutant p27, the p27 homology donor plasmid generated above was subjected to site-directed mutagenesis using NEB standard protocol and Q5 polymerase with the following forward primers:

Y88F: 5’- TTCTTCTACAGACCCCCGCGGCCCCCC -3’

Y89F: 5’- TTCTACTTCAGACCCCCGCGGCCCCCC -3’

and reverse primers:

Y88F: 5’- GTAGAAGAACTCGGGCAAGCTGCCCTT-3’

Y89F: 5’- GAAGTAGAACTCGGGCAAGCTGCCCTT-3’

All plasmids were checked by Sanger sequencing to check for introduction of the mutation and absence of any other mutations.

To generate endogenously-tagged p27-GFP cells, either p27WT or p27mutant, the pX330 p27 gRNA plasmid and the p27 homology donor plasmid generated above were transfected into hTert-RPE1 mRuby-PCNA cells^33^ at a ratio of 1:1 using Lipofectamine 2000, according to the manufacturer’s instructions (Invitrogen). Cells were allowed to expand for seven days before FACS sorting to select GFP positive cells. Individual GFP expressing cells were sorted into 96-well plates containing 50:50 conditioned:fresh growth media. Conditioned growth media had been pre-filtered through a 0.2μm filter to remove any cell debris. Clones were left to expand for 10-14 days and visualised on the Opera high-throughput microscope (PerkinElmer) to determine which clones had p27-GFP expression. Candidate clones were expanded and further validated.

#### p27 Western blotting

Samples were directly lysed by addition of Laemmli buffer. Whole cell lysates were loaded onto 4-20% Tris-Glycine Novex precast gels (ThermoFisher) followed by transfer to PVDF membranes. After protein transfer, membranes were blocked in either Blocking Buffer (5% milk in TBS with 10% glycerol) or ROTI-Block (for phospho-specific antibodies). Membranes were blocked for 1hr at RT with rocking before primary antibody diluted in Blocking Buffer was added and membranes were incubated overnight at 4°C with rocking. Membranes were washed three times in TBS+0.05%TritonX-100 (TBS/T) and anti-mouse or anti-rabbit HRP-conjugated secondary antibodies (CST) were diluted 1:2000 in Blocking Buffer or ROTI-Block and incubated with membranes for 1 hr at RT, with rocking. Membranes were washed three times in TBS/T and visualised using Clarity Western ECL Substrate (Bio-Rad). All blots were scanned on the Azure c300 imaging system. Blot quantification was performed in ImageJ. Antibodies used for Western blotting in this study are: p27 (BD, ms, 1:1000), vinculin (Sigma, V4505, ms, 1:2000), CDK2 (Santa Cruz, sc163, rb, 1:500), CDK4 (CST 12790, rb 1:1000), GAPDH (Santa Cruz sc-47724; 1:2000) Src (CST 2109 1:500) and Abl (CST 2862, 1:500).

#### p27 Co-immunoprecipitation

Immunoprecipitation of p27-GFP was carried out using the GFP-Trap Magnetic Agarose kit (Chromotek). Briefly, hTert-RPE1 cell lines were trypsinised and cell pellets washed in ice-cold PBS. Cells were lysed in Lysis buffer (10 mM Tris/Cl pH 7.5, 150 mM NaCl, 0.5 mM EDTA, 0.5% Igepal plus protease inhibitors) and passed 10x through a 26G needle. Lysates were centrifuged at 14,000xg for 15 min at 4’C and supernatants used for the immunoprecipitation reaction. 10% of the lysate was taken as Input fraction. The remainder was incubated with 25 uL of bead slurry, that had been pre-equilibrated in lysis buffer, for 1hr, with rotation, at 4’C. Supernatant was removed, beads were washed three times in Wash buffer (10 mM Tris/Cl pH 7.5, 150 mM NaCl, 0.5 mM EDTA) and protein eluted by boiling in Laemmli buffer for 10 min. All of the eluate was loaded onto a gel for analysis by western blot.

#### Growth curves

Cells were plated at a density of 1 x104 per well of a 6 well tissue culture plate and four fields of view per well were imaged using the 4x objective, every 6 hr for five days using the Incucyte (Essen Biosciences). Each experiment was performed in triplicate and results represent the mean and standard deviation of three measurements. Cell growth was calculated automatically as cell confluency change over time using the Incucyte custom software.

#### Live cell imaging

Live cell imaging was performed on the Opera HC spinning disk confocal microscope (PerkinElmer), with atmospheric control to maintain cells at 37°C, 5% CO2 and 80% humidity. Cells were plated on 384 well CellCarrier (PerkinElmer) plates at a density of 1000 cells/well 24 hr prior to imaging. Cells were imaged using a 20X (N.A. 0.45) objective at 10 min intervals for 24-72 hr in Phenol-Red Free DMEM + 10% FBS and 1% P/S.

#### Kinase Inhibitors

Saracatinib and Nilotinib were purchased from SelleckChem, reconstituted in DMSO and stored in aliquots at -20°C.

#### siRNA transfection

Cells were transfected 48 hr prior to analysis. Cells were transfected with 20 nM final concentration of siRNA using Lipofectamine RNAiMax, according to the manufacturer’s protocol (Invitrogen). Briefly, per well of a 384 well plate, 40 nl of Lipofectamine RNAiMAX was mixed with siRNA in 10 µl OPTIMEM. To this, 20 µl cells in media, at a density of 5 x 10^4^ cells/mL was added, and cells were incubated at 37°C. siRNAs used in this study were OnTARGETplus pools (Dharmacon) targeting SRC and ABL1 and a custom siRNA targeting CDKN1B CAAGUGGAAUUUCGAUUUUtt (Ambion Silencer Select s2837).

#### Immunostaining

Cells were fixed in an equal volume of media to warm 8% formaldehyde in PBS for 5 min at 37°C. Cells were permeabilised in PBS/0.2% TritonX-100 for 5 min at RT and then blocked in 2% BSA in PBS (Blocking buffer) for 1 hr at RT. Cells were incubated in primary antibody diluted in Blocking buffer for either 2 hr at RT or overnight at 4°C. Cells were washed three times in PBS and then incubated with Alexa-conjugated secondary antibodies (ThermoFisher) diluted in Blocking Buffer for 1 hr at RT. Cells were washed three times in PBS and incubated in 1 µg/mL Hoechst 33258 (Sigma) diluted in PBS for 15 min at RT. Cells were imaged on the Opera HC spinning disk confocal microscope (PerkinElmer). All imaging data was uploaded to the Columbus image analysis database (PerkinElmer) for visualisation and analysis. Primary antibodies used for immunostaining in this study are: Phospho-Ser807/811 Rb (CST 8516 1:2000), p27 (CST 3688 1:1000) and Src (CST 2109 1:1000).

#### Automated image analysis of fixed cells

All quantitative image analysis on fixed cells was performed using Columbus software (PerkinElmer) following the protocol described below.

*Measuring nuclear intensities of proteins:* Nuclei were segmented based on Hoechst intensity. Nuclei at the edge of the image and nuclei < 100 µm^2^ were excluded. Mitotic nuclei were also excluded by excluding nuclei with Hoechst intensity above a defined threshold (that was defined individually for each experiment based on Hoechst staining intensity and imaging conditions). The fluorescence intensity of individual proteins was then calculated in each interphase nucleus. To calculate the fraction of cells in G0 vs G1/S/G2 we defined a threshold for nuclear phospho-Rb staining. Rb hyperphosphorylation correlates with passage through the Restriction Point and CDK2: CyclinE activity^72^. Cells with a nuclear intensity of phospho-Rb below that threshold are classed as G0/quiescent and cells above that threshold as proliferating.

### Structural Homology Modeling

Homology modeling was performed to evaluate pTyr recognition in the kinase domains of Abl and Src. For Abl, the substrate peptide Abltide (PDB: 2G2F)^37^ was used as a structural reference, and the –1 position relative to the phosphorylatable tyrosine was computationally mutated to pTyr. For Src, the kinase domain (PDB: 3D7T)^38^ was structurally aligned to the Abl–Abltide complex to model substrate binding, and the +1 position was similarly mutated to pTyr. In both cases, rotamers were manually selected to minimize steric clashes. All modeling, mutation, and structure visualization steps were performed using UCSF Chimera^73^.

### Computational analysis of determinants of specificity quantifying evolutionary conservation and identifying cancer somatic mutations perturbing them

We compiled a multiple sequence alignment (MSA) of kinase domains corresponding to the catalytic part of all known human protein kinases, from which we generated a second alignment in fastA file format containing no sequence gaps. In parallel, we compiled a set of publicly available unique missense somatic cancer point mutations from COSMIC version 82^74^ and, to enable the mapping of mutants, a mutant fastA file containing both wild type and mutant counterpart versions of all coding missense variants by running Ensembl’s variant effect predictor (VEP) resource v87.18 as described^75^ using human’s genome sequence version GRCh38.p78. Subsequently, we mapped the MSA fastA file with the mutant fasta file to build a final mapping table containing all cancer point-mutations matched across the aligned human kinase domains. All computational work was coded using purpose-made python scripts.

Images representing the mapping of information into the kinome tree were illustrated using Kinome Render (KR) version 1.4^76^. The first dendrogram includes all basic residues present in Abl-like 328 or Src-like 472 positions across the MSA. To generate it, we wrote a purpose-made python script that takes the MSA fastA file (with sequence gaps) and selects kinases showing basic residues (arginine and lysine) in alignment positions 500 (corresponding to Abl^R328^) and 1208 (corresponding to Src^R472^). These parsed kinases are then annotated on the kinome tree accordingly. Similarly, we generated the second dendrogram by running a python script that uses the MSA file as well as the final mapping table obtained in the computational mapping of missense cancer point-mutants, and selects all kinases whose Abl^R328^ or Src^R472^ positions presenting basic residues have been reported in COSMIC to be mutated in cancer.

### Compilation and computational analysis of YY sites

To compile a comprehensive list of YY sites, including those reported as phosphorylated, YY sites were extracted by scanning the human reference proteome (UP000005640)^21^, and phosphorylation annotation data were retrieved from PhosphoSitePlus^22^. This enabled the construction of a background dataset comprising all possible YY motifs, from which the subset reported as phosphorylated was defined as the foreground set for comparative analyses. These two custom sets were subsequently used in g:Profiler^77^ to obtain enriched gene sets. Reported substrates of Abl and Src kinases were also retrieved from PhosphoSitePlus^22^ to enable filtering for shared YY sites.

### *In vivo* mouse modeling comparing cancer progression for different Bcr-Abl mutants

A 30 mg/mL stock solution of 5-fluorouracil (5-FU; Sigma, F6627-10G) was prepared in sterile PBS. The solution was heated on a 95 °C heat block for 5 minutes to dissolve the compound and mixed thoroughly. After cooling to 37 °C, 100 µL of the prepared solution (equivalent to a 3 mg dose, ∼150 mg/kg) was administered via intraperitoneal injection to C57BL/6J mice.

Four days after injection, 5-FU–treated mice were euthanized, and femurs and tibias were dissected using sterile techniques and equipment. Bones were cleaned and submerged in 70% ethanol for 1 minute for surface sterilization. For each pair of bones (one femur and one tibia), a 0.65 mL tube was perforated at the bottom using a 21G needle and nested inside a 1.5 mL collection tube containing 100 µL of Iscove’s Modified Dulbecco’s Medium (IMDM) containing 10% fetal bovine serum (FBS). One end of each bone was nipped close to the tip with scissors and placed inside the perforated tube. Tubes were centrifuged at 15,000 × g for 15 seconds at 4 °C. The flushed bone marrow (BM) cells were collected, and red blood cells (RBCs) were lysed by adding 1 mL of RBC lysis buffer (Sigma, R7757) to each tube. All cell suspensions were pooled into a 15 mL tube. Samples were incubated for 4 minutes at room temperature, then washed twice with IMDM (10% FBS), and finally resuspended in 1 mL of the same medium for counting. Cells were plated in 6-well plates at a density of 1 million cells/mL in a total volume of 4 mL IMDM supplemented with 10% FBS, 5% WEHI-conditioned media, 6 ng/mL IL-3, 10 ng/mL IL-6, and 100 ng/mL SCF. WEHI-conditioned media was generated by culturing WEHI cells (ATCC CRL-1702) in 45% IMDM, 45% Dulbecco’s Modified Eagle Medium (DMEM), and 10% FBS until the media turned orange. Cultures were incubated at 37 °C in a tissue culture incubator.

The procedure for retroviral transduction of bone marrow cells and transplantation into lethally irradiated recipient mice was adapted from Pear *et al*^55^. On day 5, the supernatant from cultured bone marrow cells was collected, and cells were pelleted by centrifugation at 250 × g for 4 minutes at room temperature. Pellets were resuspended in 1 mL of retroviral supernatant of choice and 3 mL of IMDM-based culture media (10% FBS, 5% WEHI-conditioned media, 6 ng/mL IL-3, 10 ng/mL IL-6, 100 ng/mL SCF) containing 4 µg/mL polybrene. This mixture was returned to the original 6-well plate containing any remaining adherent cells. Plates were centrifuged at 300 × g for 45 minutes at 37 °C and then returned to a tissue culture incubator at the same temperature. On day 6, the infection process was repeated as above. On day 7, transplant-recipient mice were gamma-irradiated with 7 Gy. Transduced bone marrow cells were collected by transferring the culture media from each well and detaching adherent cells by incubating with 1 mL PBS containing 5 mM EDTA. Detached cells were combined with non-adherent cells from the same sample. The suspension was centrifuged at 250 × g for 5 minutes, and retroviral supernatants were discarded into a container containing 50% bleach. Cells were washed twice with PBS, counted, and resuspended in PBS before analysing GFP expression by flow cytometry. Six hours after mice were gamma-irradiated, an equal number of 250,000 cells was injected into each recipient mouse via the tail vein in 100 µL of PBS.

### Histopathologic analyses

Spleens from Bcr-Abl^WT^, Bcr-Abl^R328A^ and Bcr-Abl^R328M^ endpoint mice were collected, weighed, and processed for histological examination. Samples were fixed, embedded, sectioned, and stained with haematoxylin and eosin (H&E) to assess disease progression and tissue morphology, and with an anti-Ki-67 antibody (ab16667, 1:1000 dilution) for immunohistochemistry (IHC) to evaluate Bcr-Abl–driven cellular proliferation. Slides were scanned using the Aperio slide scanner (Leica Biosystems) and Ki-67–stained IHC images were segmented into uniform tiles of 10^5^ µm² using Halo software (v3.3, Indica Labs). Analysis focused on the spleen region, and a trained HALO tissue classifier was used to exclude background and areas with incomplete staining. Stain vectors for haematoxylin and DAB were determined via the colour deconvolution algorithm in QuPath (v 0.5.0). The modified CytoNuclear algorithm (v2.0.9) in Halo was then applied to identify Ki-67–positive cells and calculate the immunopositive rate, defined as the proportion of positively stained cells relative to the total cell count within each tile. To ensure accuracy, all spleen samples were independently assessed in a blinded manner by two immunologists.

### Staining of HSCs and progenitor cells

Bone marrow (BM) cells were harvested and red blood cells lysed as described above. Lineage positive cells were depleted using mouse lineage cell depletion kit (Miltenyi Biotech) and the following antibodies were used for staining: Rat monoclonal APC anti-mouse CD117, BD Biosciences, Cat#561074; RRID: AB_10563203, Rat monoclonal APC/Cy7 anti-mouse CD117, BioLegend, Cat#105826, RRID: AB_1626280, Rat monoclonal PE/Cy7 anti-mouse Ly-6A/E, BioLegend, Cat#108114; RRID: AB_493596; Armenian hamster monoclonal BV421 anti-mouse CD48, BioLegend, Cat#103427; RRID: AB_10895922, Rat monoclonal PE anti-mouse CD150, BD Biosciences, Cat#562651; RRID: AB_2737705, Rat monoclonal PE anti-mouse CD16/32, BioLegend Cat#101307, RRID: AB_312806, Rat monoclonal PE anti-mouse CD127, BioLegend Cat#135009, RRID: AB_1937252, Rat monoclonal APC anti-mouse CD34, BioLegend Cat#119309, RRID: AB_1236482. Cells were analyzed using an Aria cell sorter (Becton Dickinson). For all flow cytometry experiments, FlowJo was used for analysis.

### Statistical Analysis

All statistical analyses were performed using GraphPad Prism (version 10.2.0) and R (v4.4.0). One-way ANOVA or unpaired two-tailed Student’s *t*-tests were applied as appropriate, with the number of biological replicates (N) and *P*-values indicated in the figure legends. Data are presented as mean ± standard deviation (SD) or standard error of the mean (SEM), as specified. Kaplan–Meier survival curves for overall survival in mice were generated using GraphPad Prism, and statistical significance between groups was assessed using the log-rank (Mantel–Cox) test.

---

**Figure S1.**
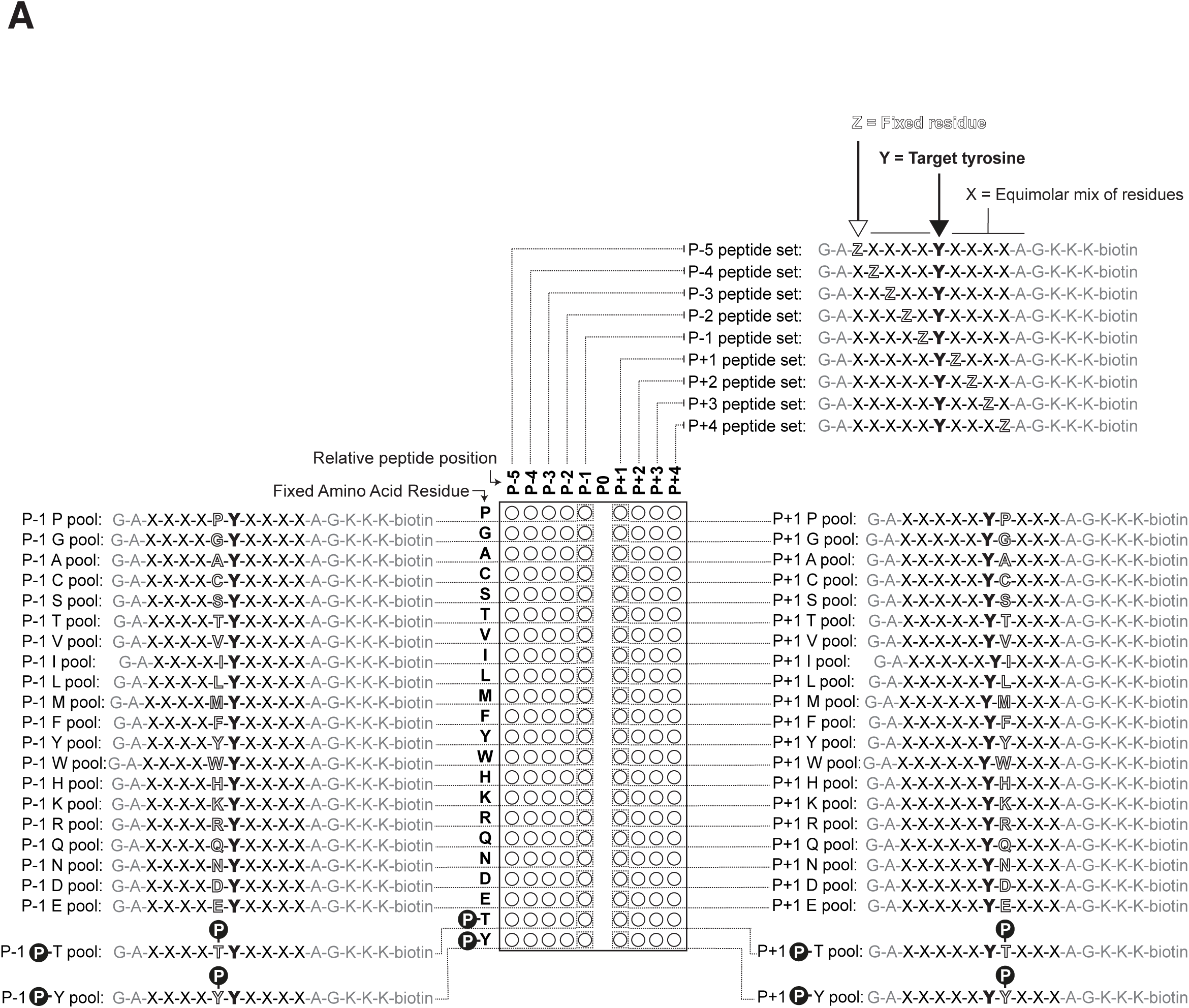
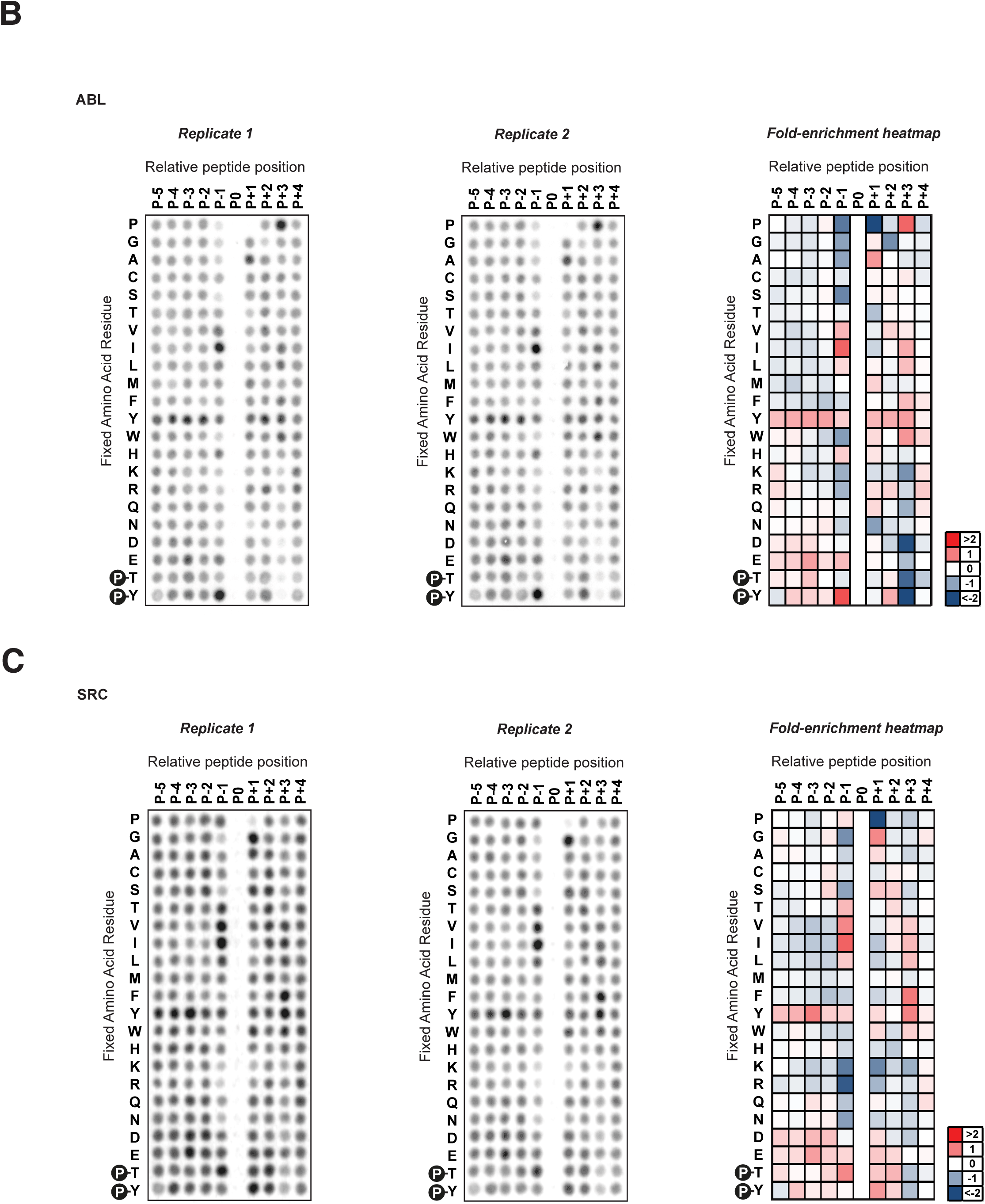
(A) Schematic representation of the structure for the different peptide pools that form a Positional Scanning Peptide Library (PSPL) screen with the target tyrosine residue that will be phosphorylated always present centrally and surrounded by an equimolar mix of other 21 residues except for one position, position Z that contains a fixed residue. In each column a different Z position is fixed from five positions N-terminal to 4 positions C-terminal from the target tyrosine, while in each row a different amino acid residue is fixed. Our screens included pre-phosphorylated phosphothreonine and pTyr residues as the two last rows. (B) Positional Scanning Peptide Library (PSPL) screening results for Abl kinases including two replicates (left and center) and column-normalized log2 fold-change enrichment heatmaps from averaged data from the two replicates (right). (C) Positional Scanning Peptide Library (PSPL) screening results for Src kinases including two replicates (left and center) and column-normalized log2 fold-change enrichment heatmaps from averaged data from the two replicates (right).

**Figure S2.**
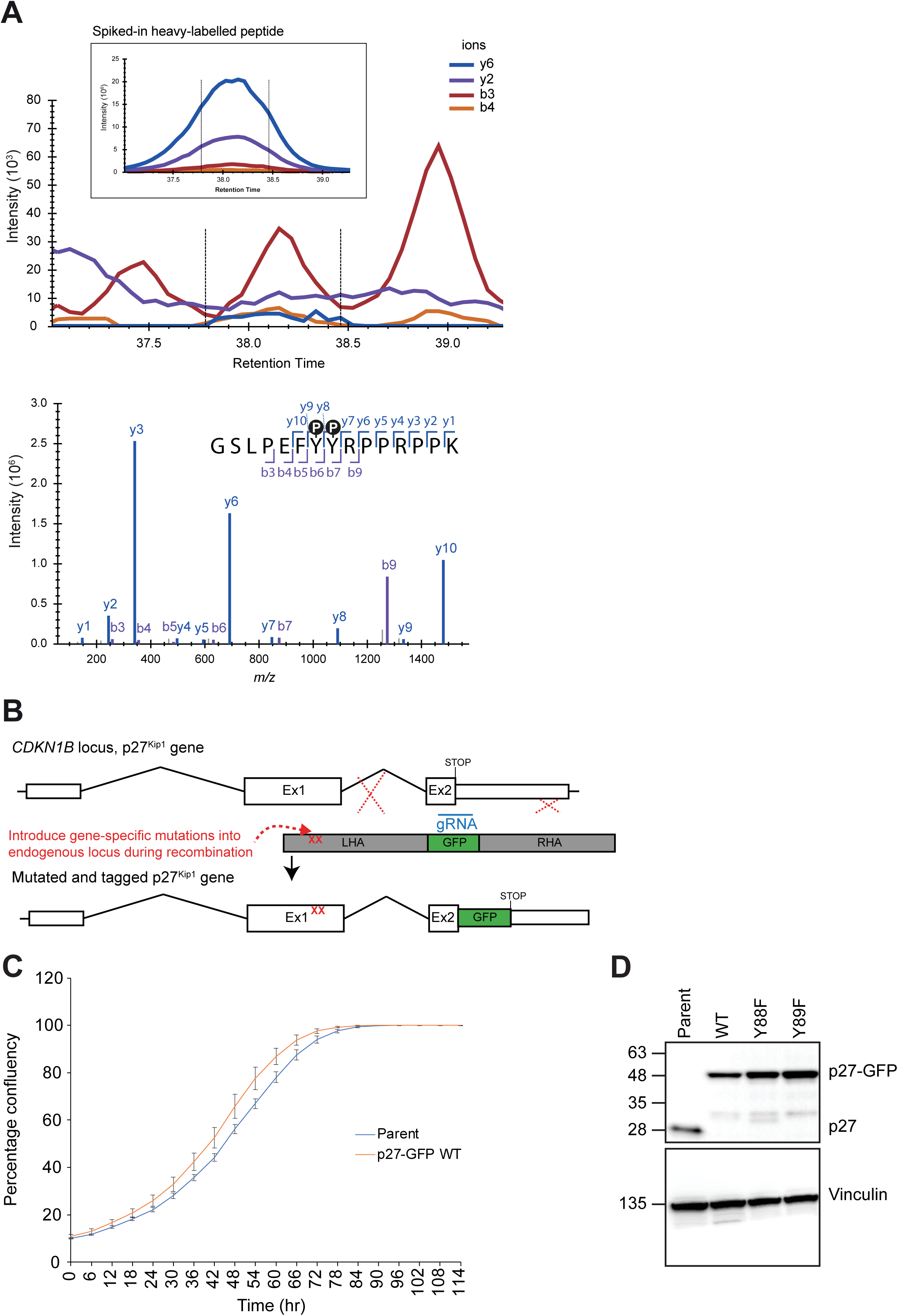
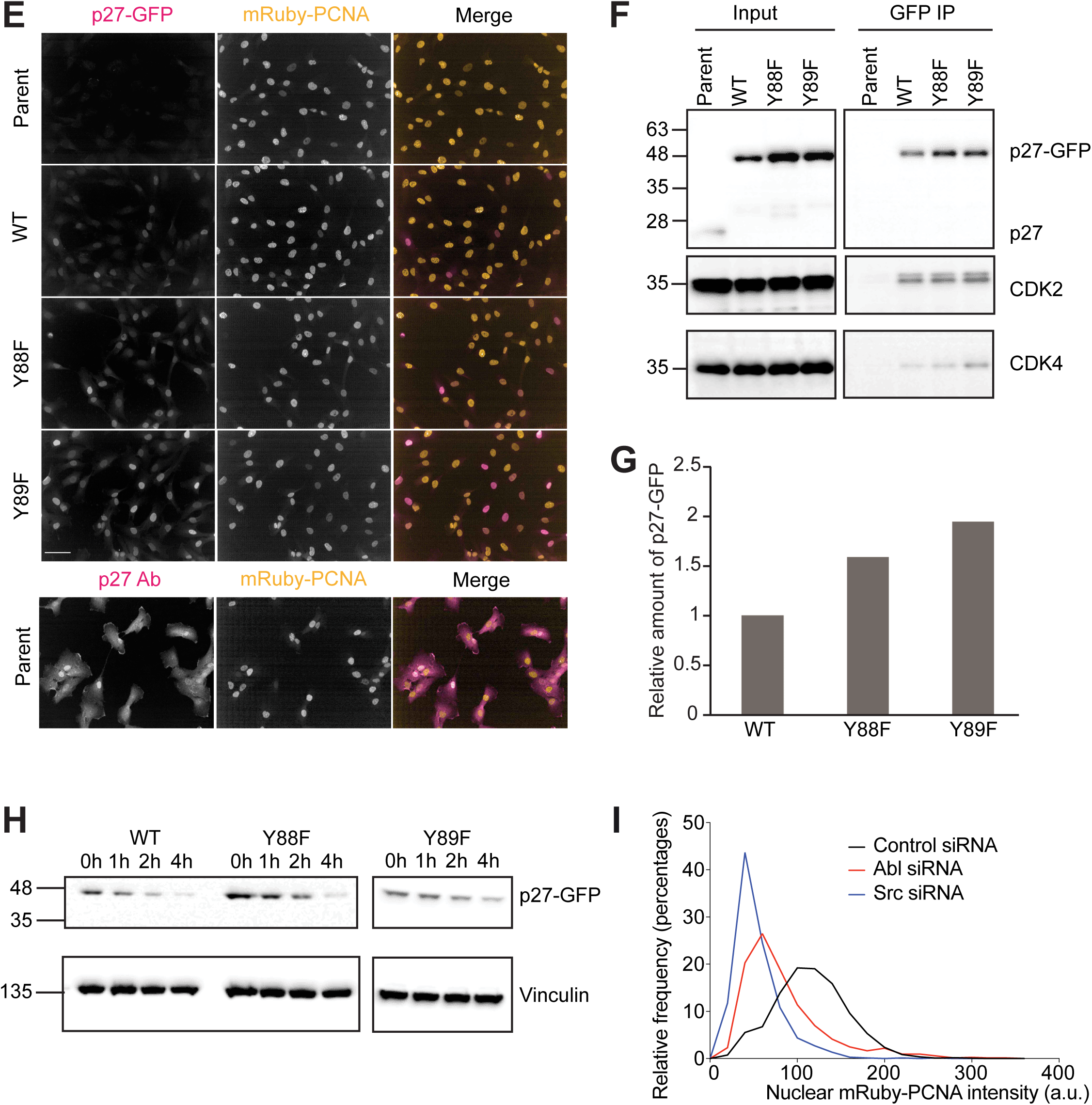
(A) Top. Extracted Ion Chromatogram for the synthetic doubly phosphorylated peptide, both in heavy-labelled spike-in and endogenous form, confirming their highly similar elution profiles and retention time. Bottom. Targeted-MS2 spectrum of the doubly phosphorylated peptide, indicating robust b- and y-ion coverage of the peptide, including the twin pTyr-pTyr residues (b6, b7 and y8, y9 respectively), supporting the empirical observation of this peptide in LC-MS. (B) Design of gene-targeting strategy to introduce a GFP tag into the *CDKN1B* locus (encoding p27) and to mutate tyrosines Y88 and Y89 to phenylalanine residues. (C) Graph showing growth curves of parental line versus p27^WT^-GFP. Mean +/- STD of n=3 is shown. (D) Western blot showing homozygous tagging of p27 to generate p27-GFP. The parental line (hTert-RPE1 mRuby-PCNA only) is shown for comparison. Vinculin is used as a loading control. (E) Four top row panels. Images showing p27-GFP (magenta in merged image) and mRuby-PCNA expression (gold in merged image) in parent, p27^WT^-GFP and p27^mutant^-GFP cell lines. Bottom row panels. Immunostaining of parent cell line with anti-p27 antibody. Both the anti-p27 antibody and the GFP tag show the same nuclear and cytoplasmic localisation of p27 protein. Scale bars are 50 μm. (F) Western blot showing that both p27^WT^-GFP and p27^mut^-GFP proteins still bind to CDK2 and CDK4. GFP immunoprecipitations of tagged proteins were performed on whole cell lysates and the eluted complexes were run on a western blot and probed for the proteins shown. (G) Graph shows quantification of p27-GFP levels in the p27^Y88F^-GFP and p27^Y89F^-GFP mutant lines compared to p27^WT^-GFP. (H) Western blots showing stability of p27-GFP proteins after 100 μg/mL cycloheximide treatment. Vinculin is shown as a loading control. (I) Quantification of nuclear mRuby-PCNA fluorescence intensity in individual cells after control, Abl or Src siRNA depletion. Low levels of PCNA expression are an indicator of quiescence since PCNA is an E2F target, activated after passing the restriction point. (Control: n=1051 cells, Abl: n=729 cells, Src n=664 cells).

**Figure S3.**
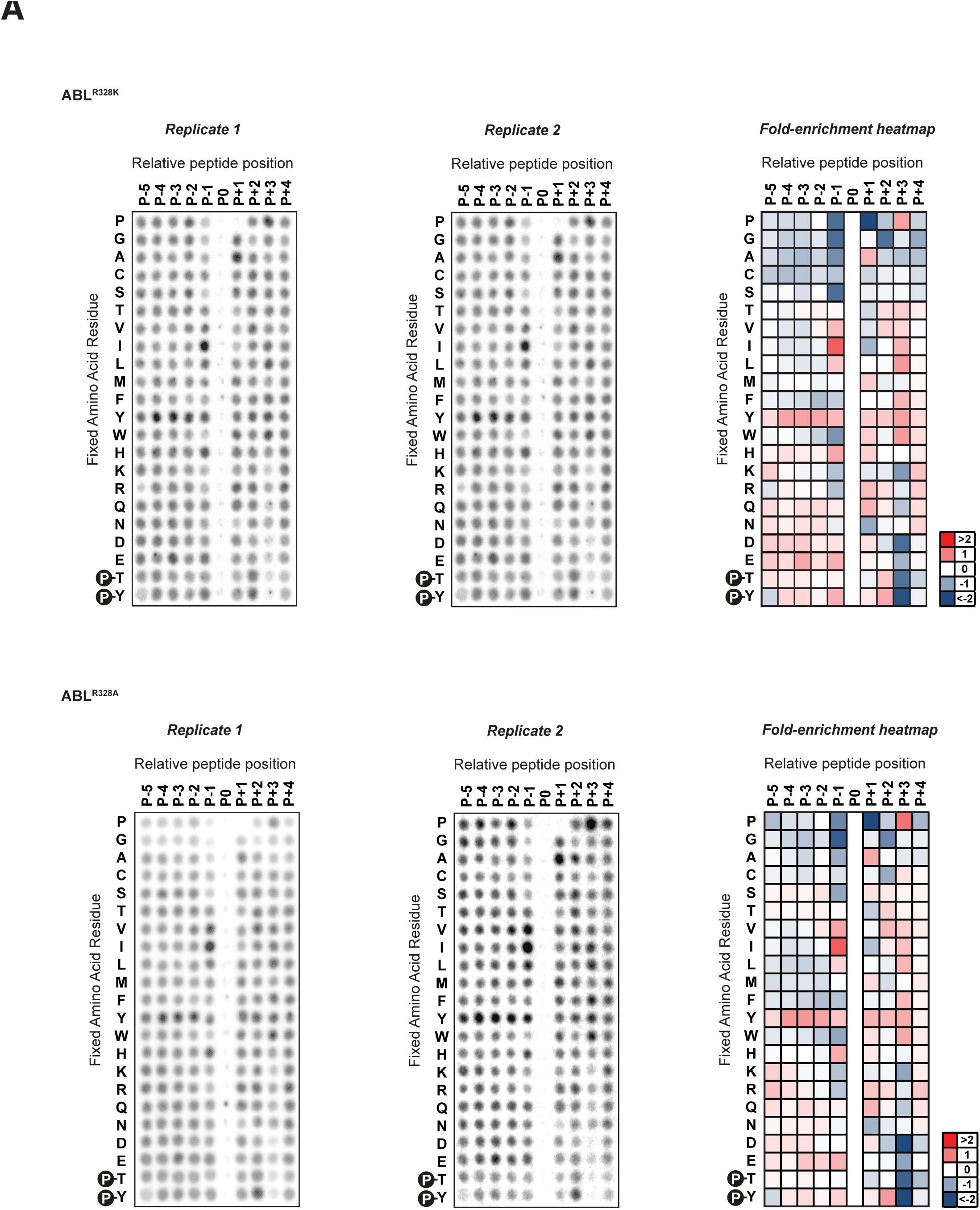
(A) Positional Scanning Peptide Library (PSPL) screening results for Abl^R328K^ (top) and Abl^R328A^ (bottom) kinases including two replicates (left and center) and column-normalized log2 fold-change enrichment heatmaps from averaged data from the two replicates (right).

**Figure S4.**
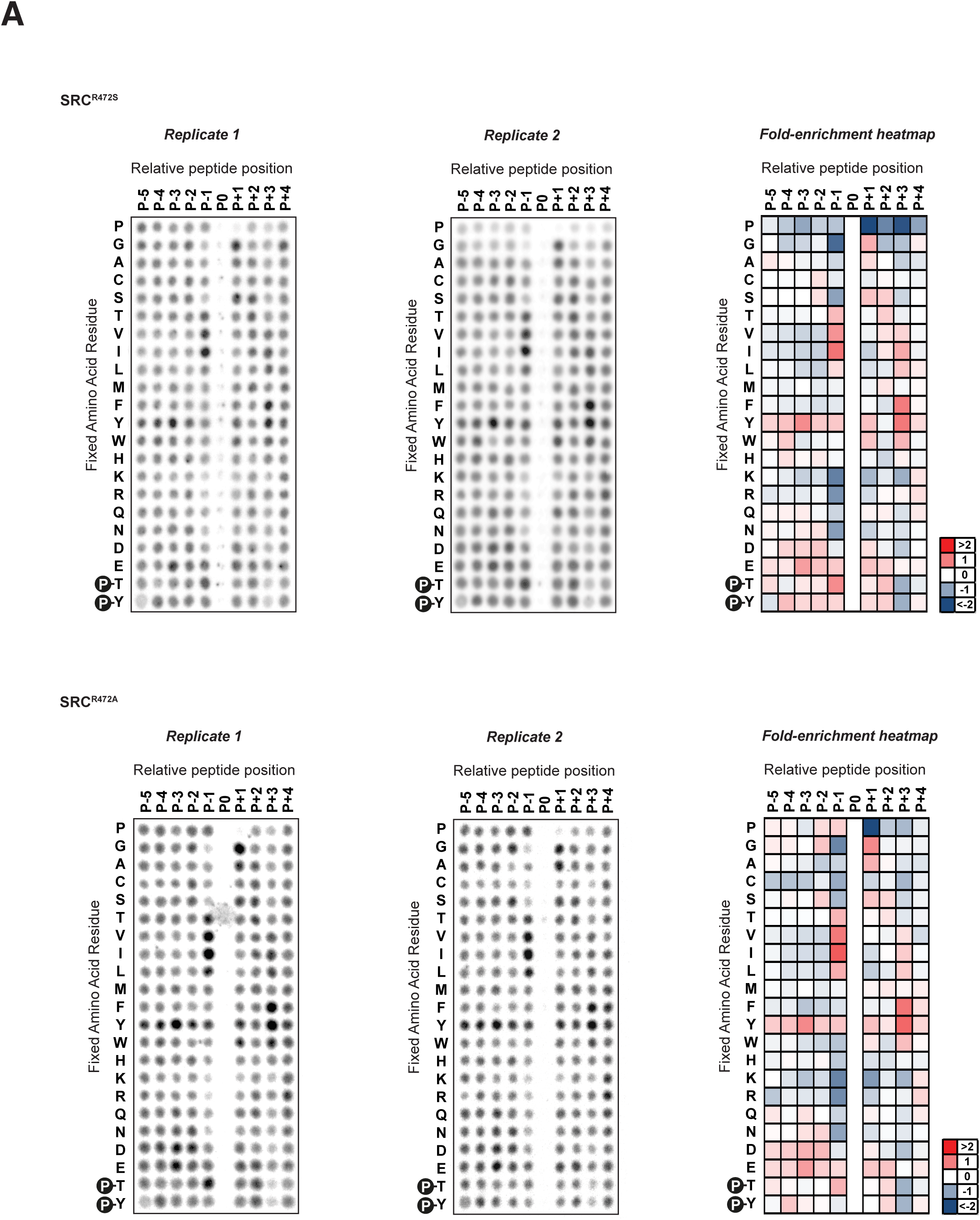
(A) Positional Scanning Peptide Library (PSPL) screening results for Src^R472S^ (top) and Src^R472A^ (bottom) kinases including two replicates (left and center) and column-normalized log2 fold-change enrichment heatmaps from averaged data from the two replicates (right).

**Figure S5.**
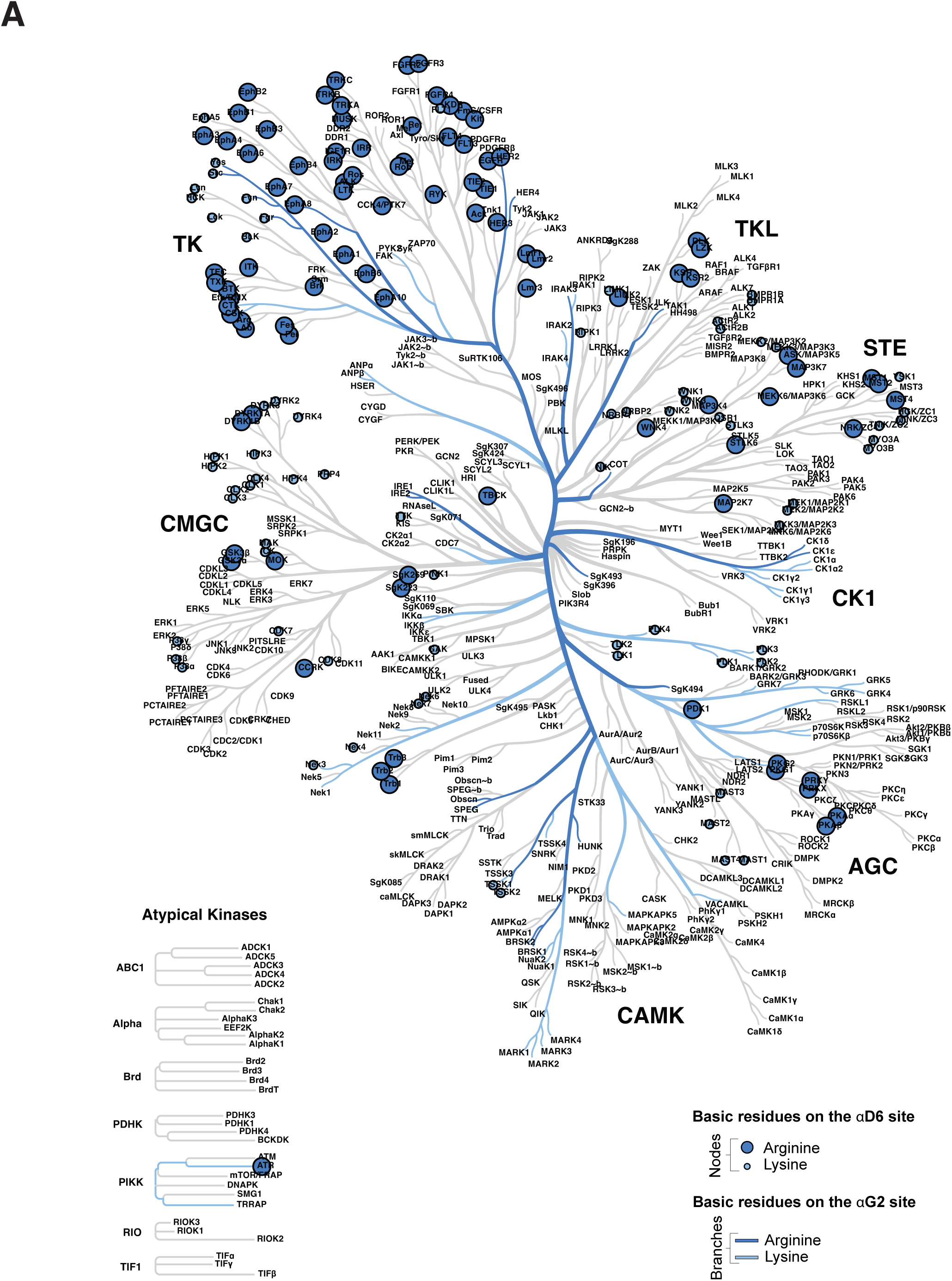
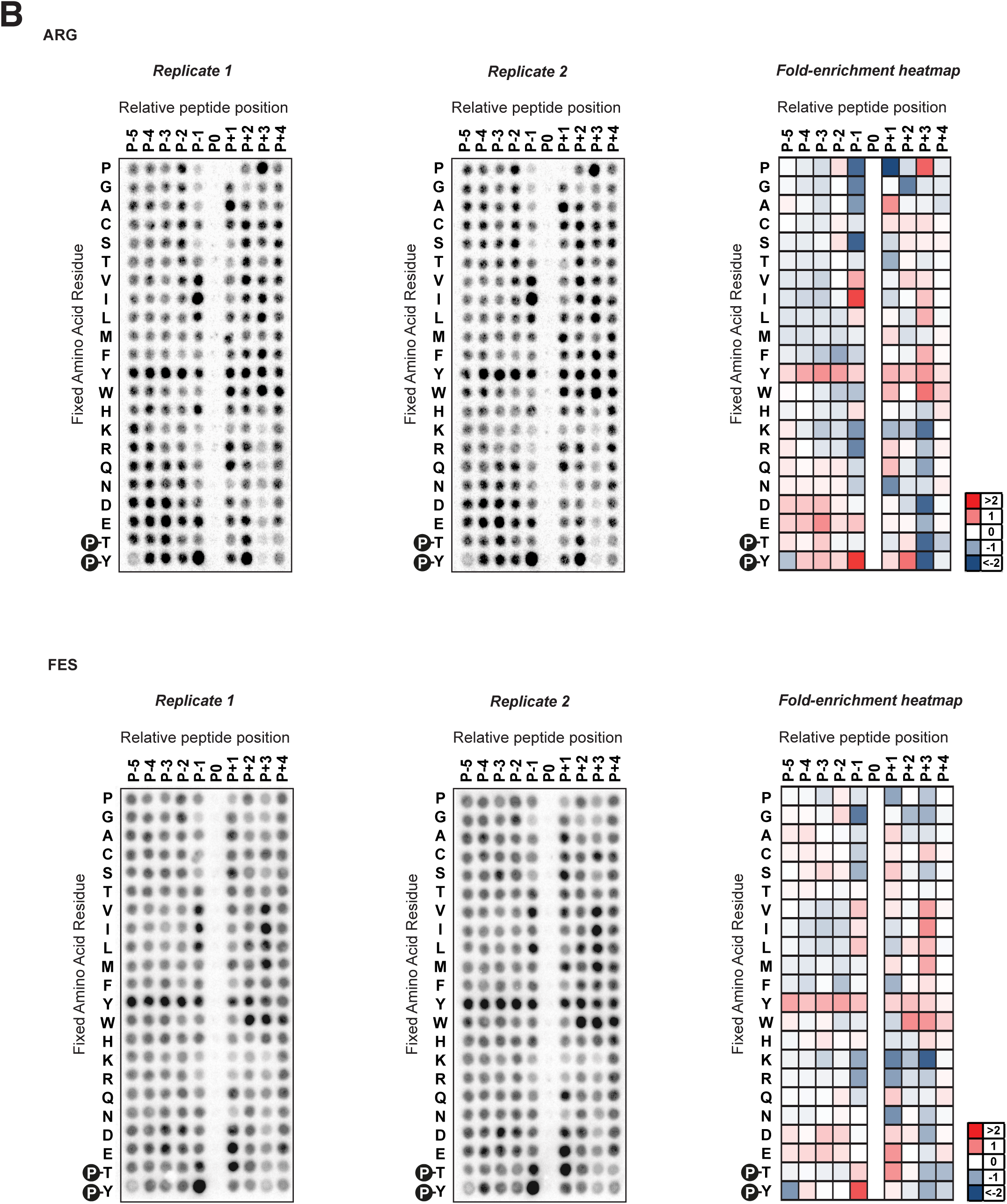
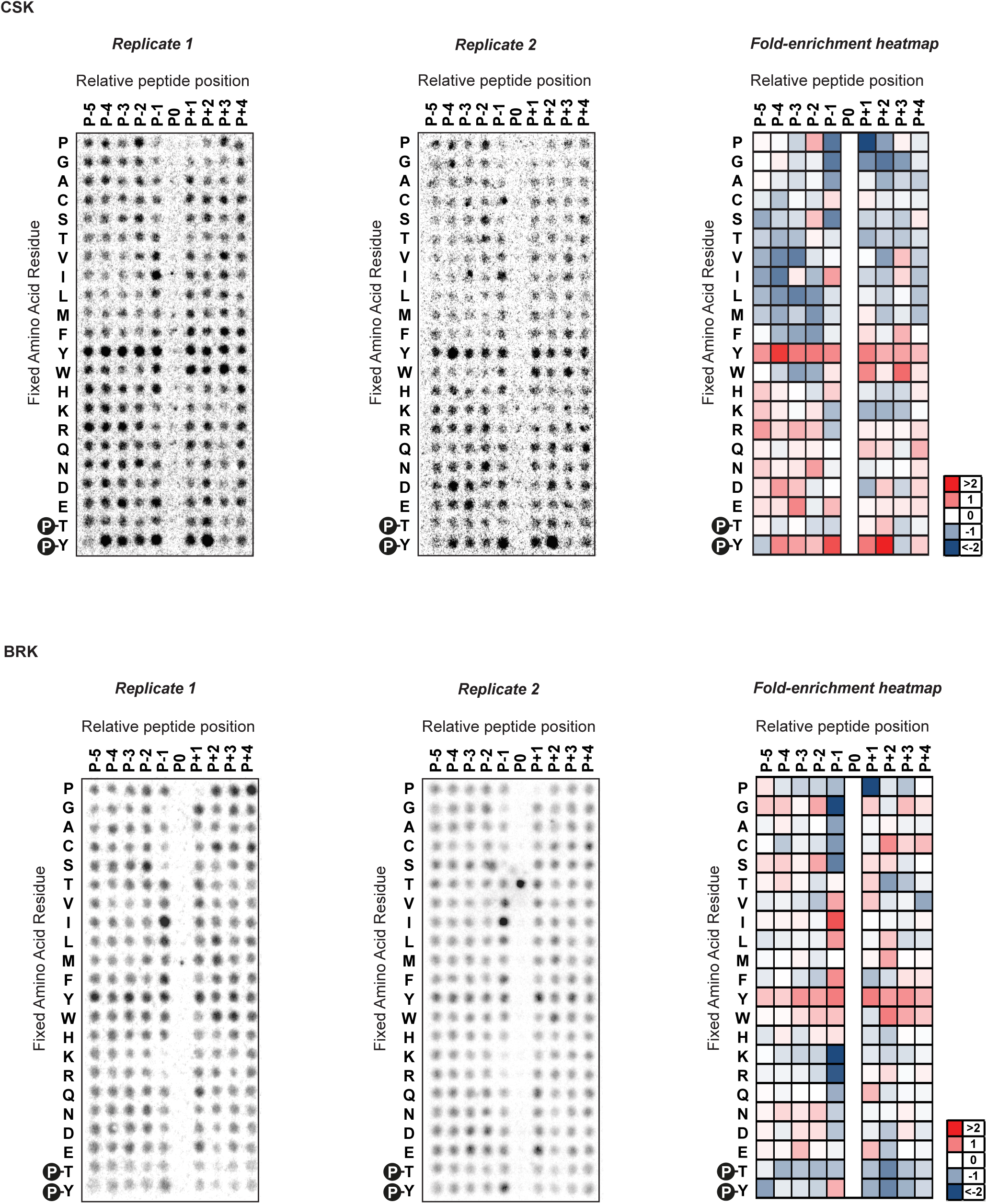
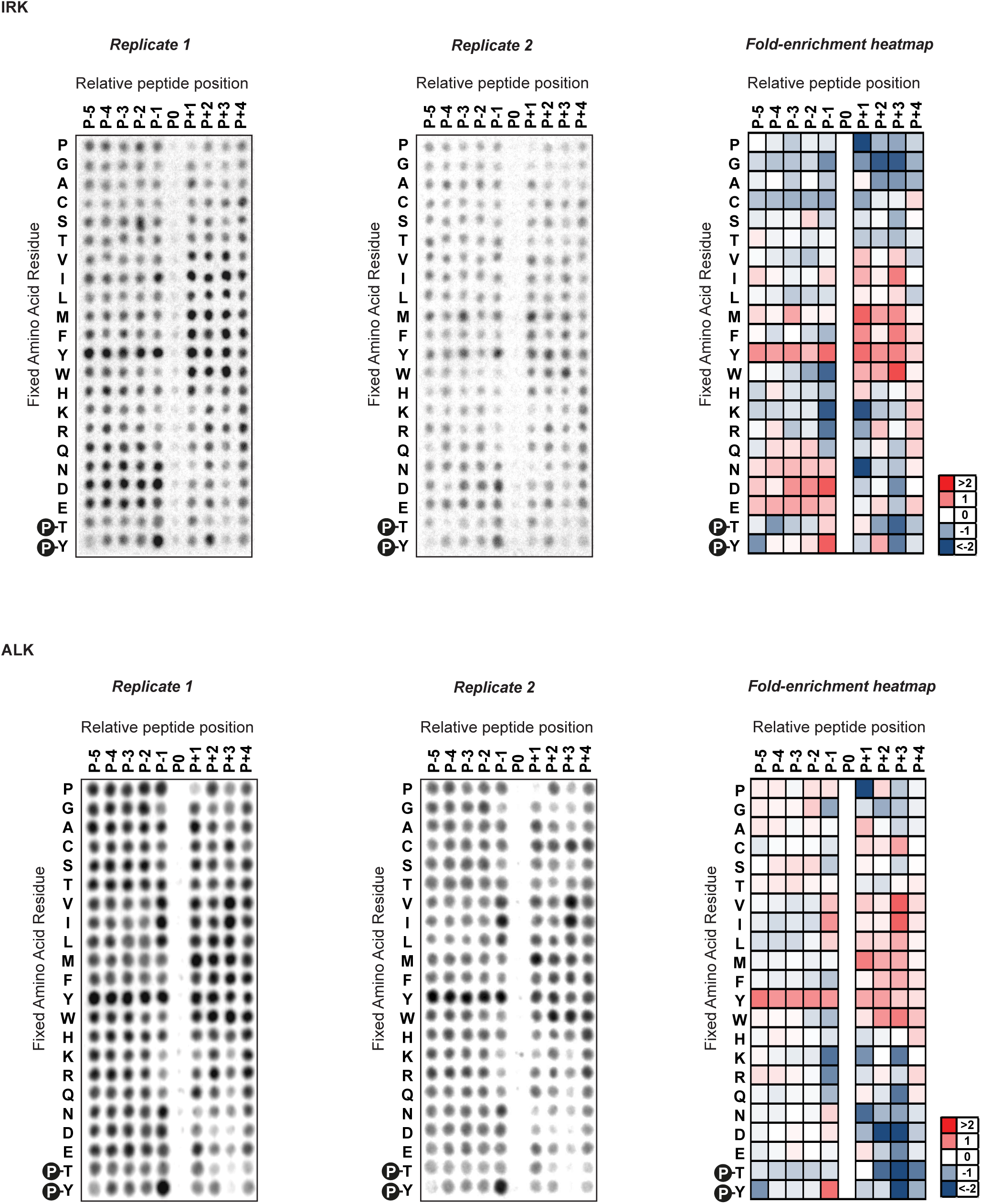
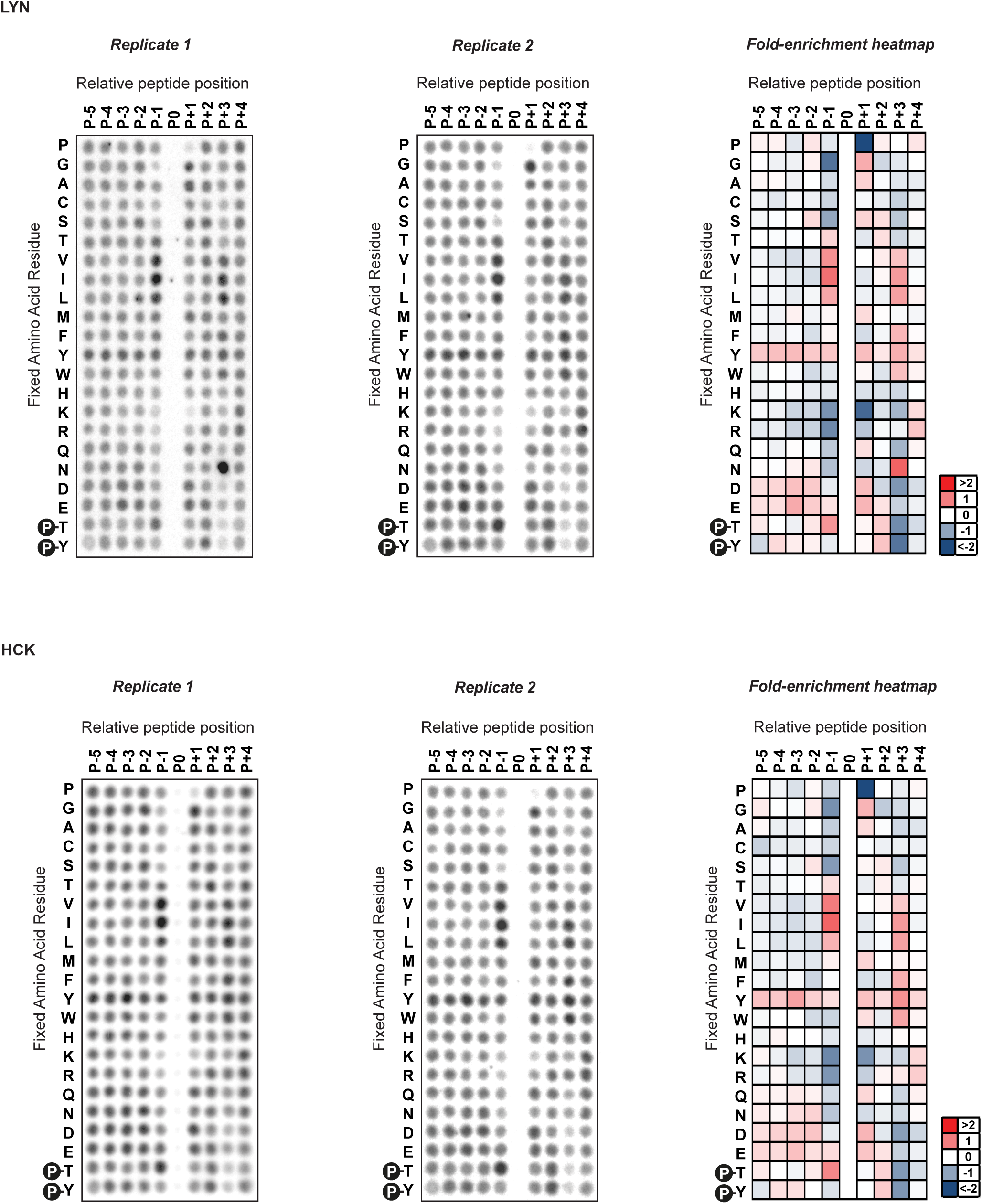
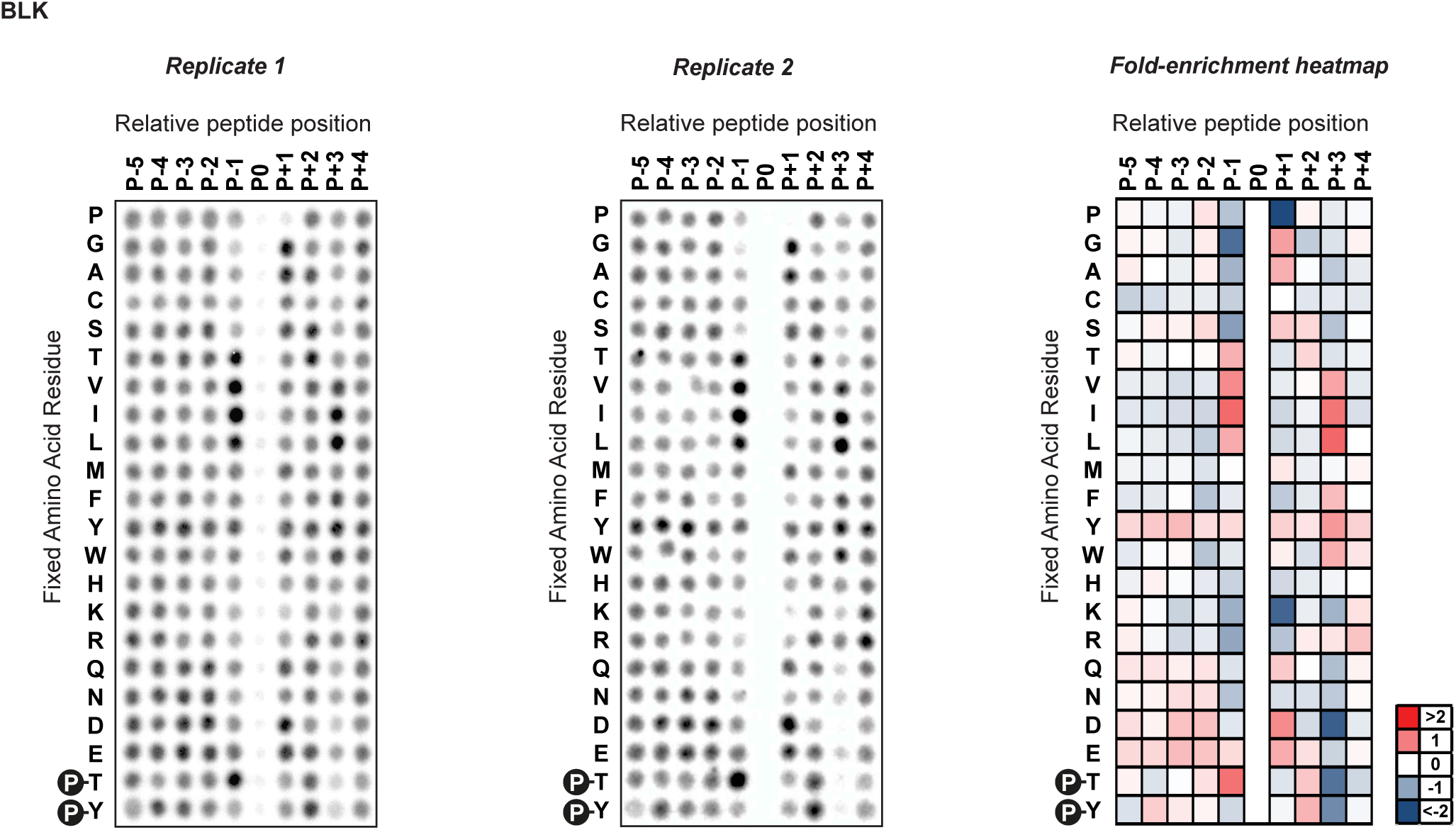
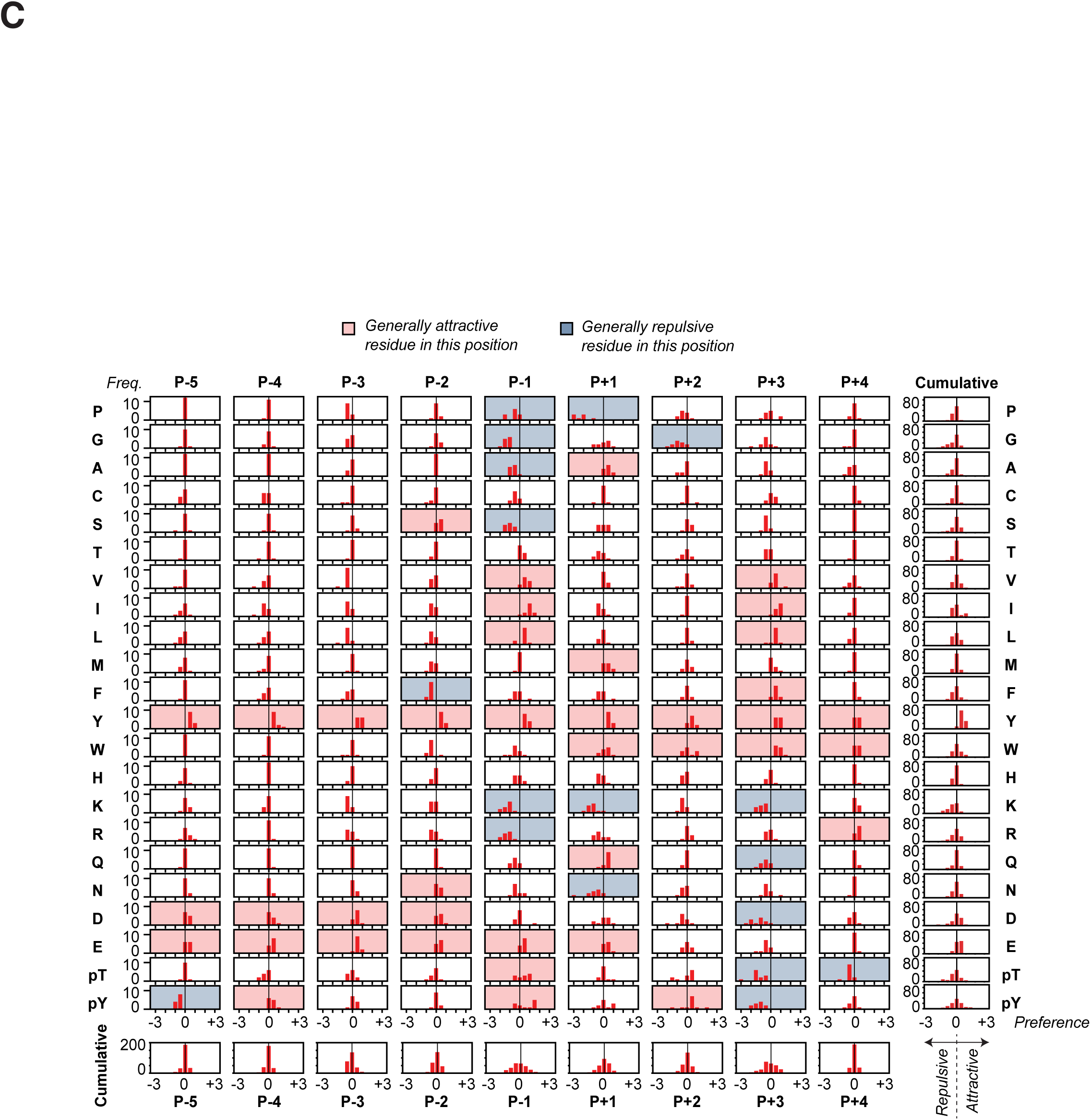
(A) Dendrogram of the phylogenetic relationship of kinases within the human proteome with branches coloured according to the identity of their *α*G2 site residue and leaves coloured according to the identity of their *α*D6 site residue with Lysine and Arginine shown in lighter-to-darker blue. (B) Positional Scanning Peptide Library (PSPL) screening results for Arg, Fes, Csk, Brk, IRK, Alk, Lyn, Hck and Blk kinases including two replicates (left and center) and column-normalized log2 fold-change enrichment heatmaps from averaged data from the two replicates (right). (C) Distributions of the favorability of the different amino acid residues in the different positions relative to fixed tyrosine phosphoacceptor in all the PSPL screens across all 11 wild-type kinases in our study, with generally favorable residues in a position coloured with red background and generally disfavorable residues in a position coloured with blue background.

**Figure S6.**
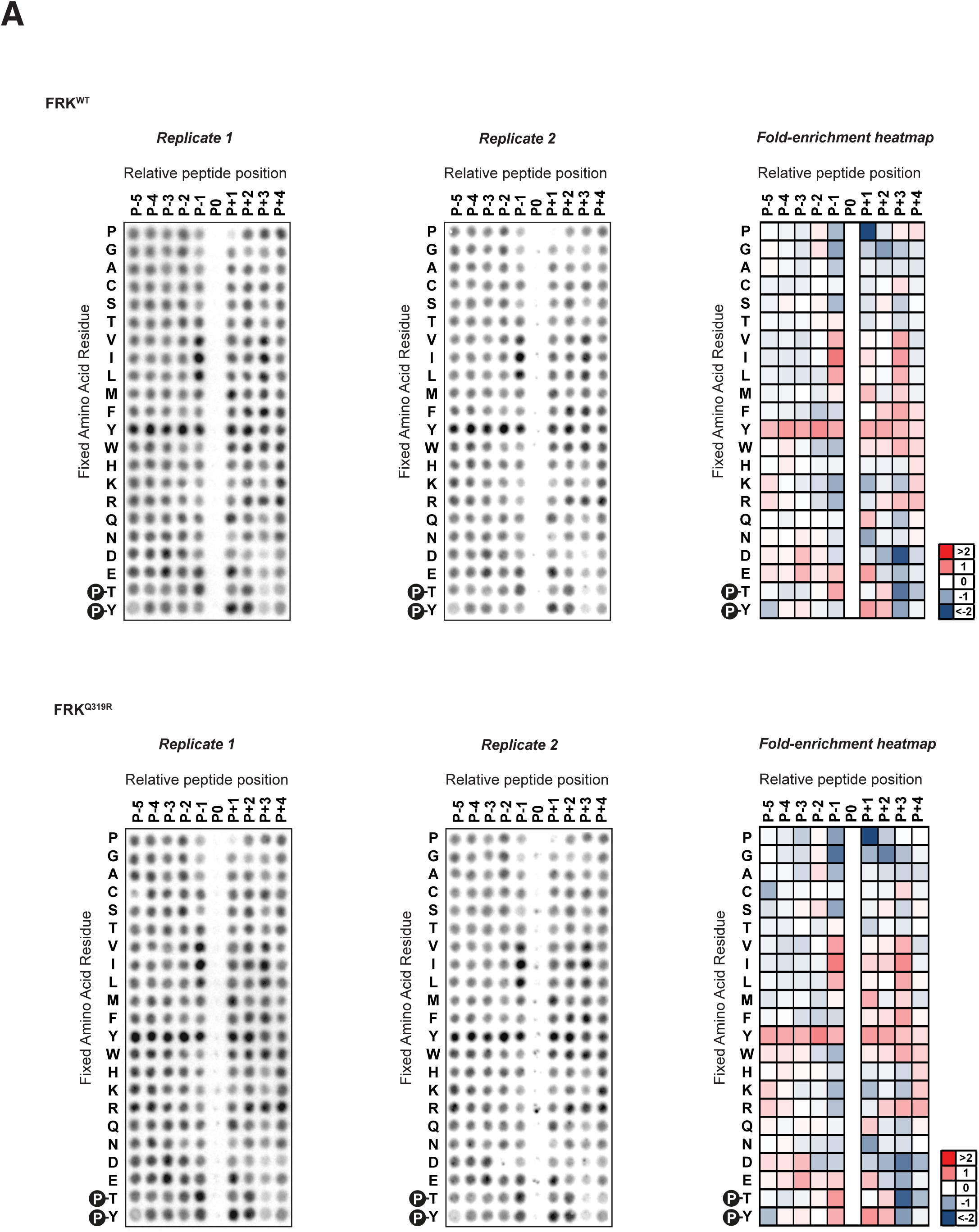
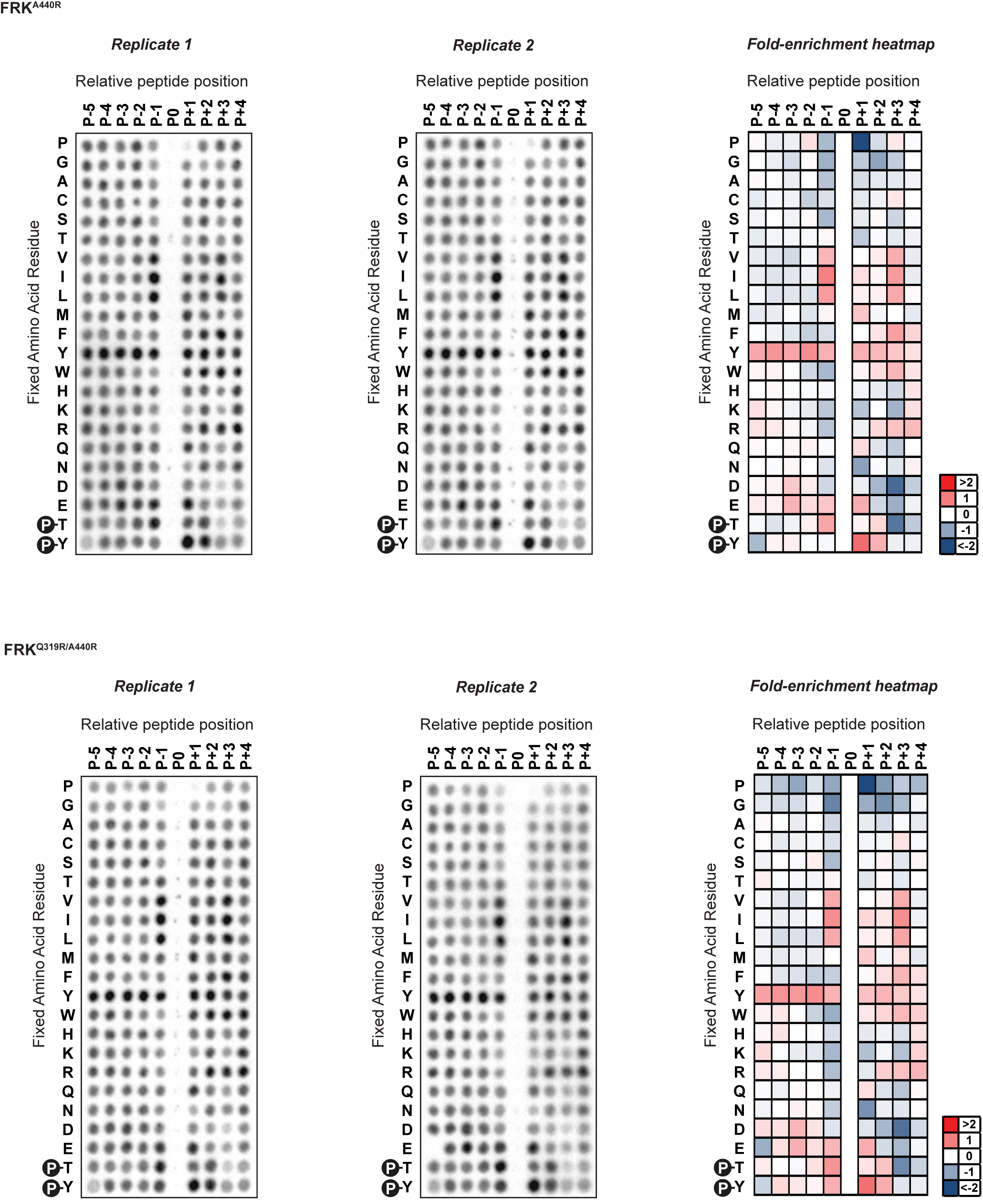
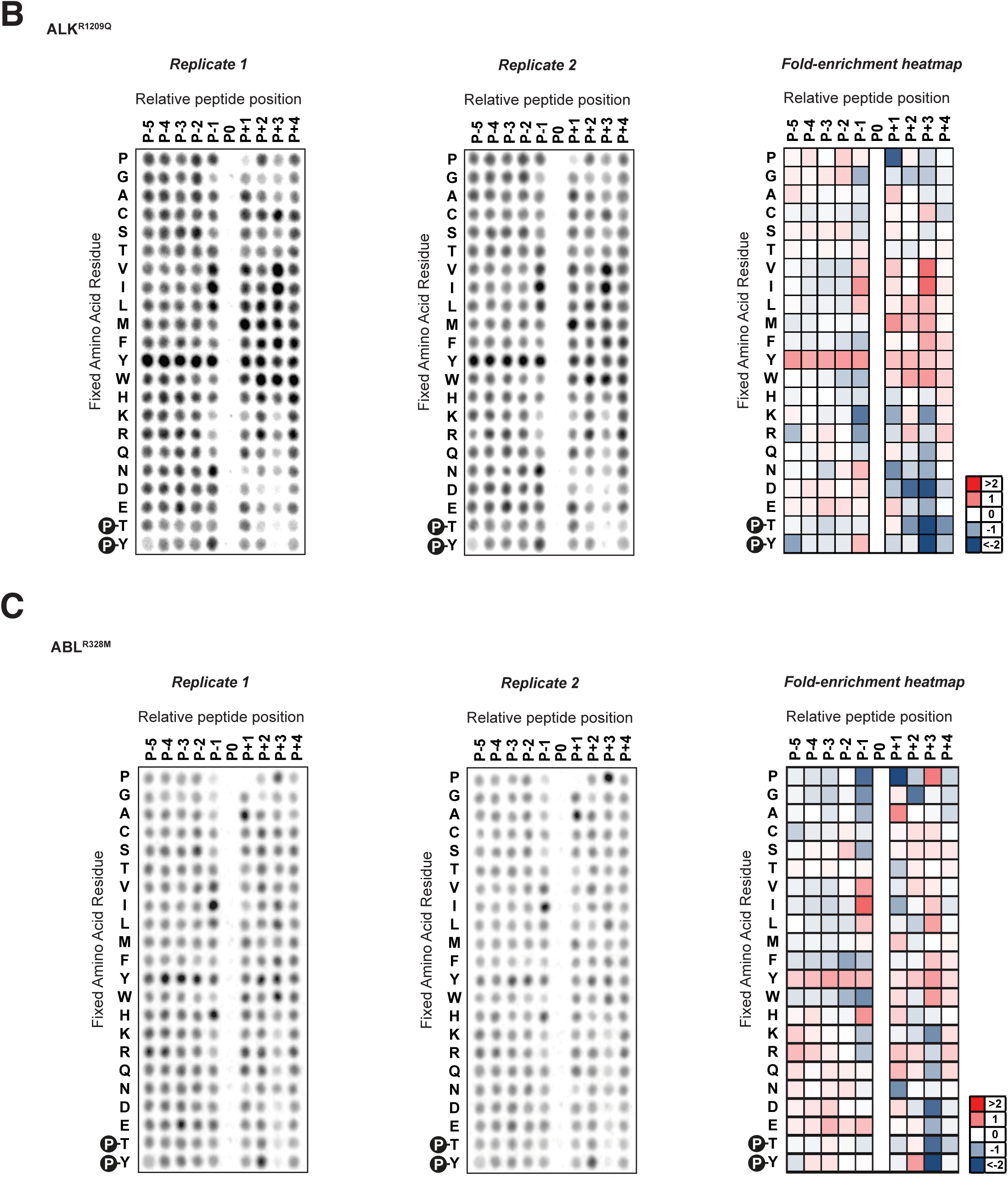
(A) Positional Scanning Peptide Library (PSPL) screening results for Frk^wt^, the *α*D6 site mutant Frk^Q319R^, the *α*G2 site mutant Frk^A440R^, and the double mutant Frk^Q319R A440R^ kinases including two replicates (left and center) and column-normalized log2 fold-change enrichment heatmaps from averaged data from the two replicates (right). (B) Positional Scanning Peptide Library (PSPL) screening results for the cancer somatic mutations Alk^R1209Q^ kinases including two replicates (left and center) and column-normalized log2 fold-change enrichment heatmaps from averaged data from the two replicates (right). (C) Positional Scanning Peptide Library (PSPL) screening results for the cancer somatic mutations Abl^R328M^ kinases including two replicates (left and center) and column-normalized log2 fold-change enrichment heatmaps from averaged data from the two replicates (right).

**Figure S7.**
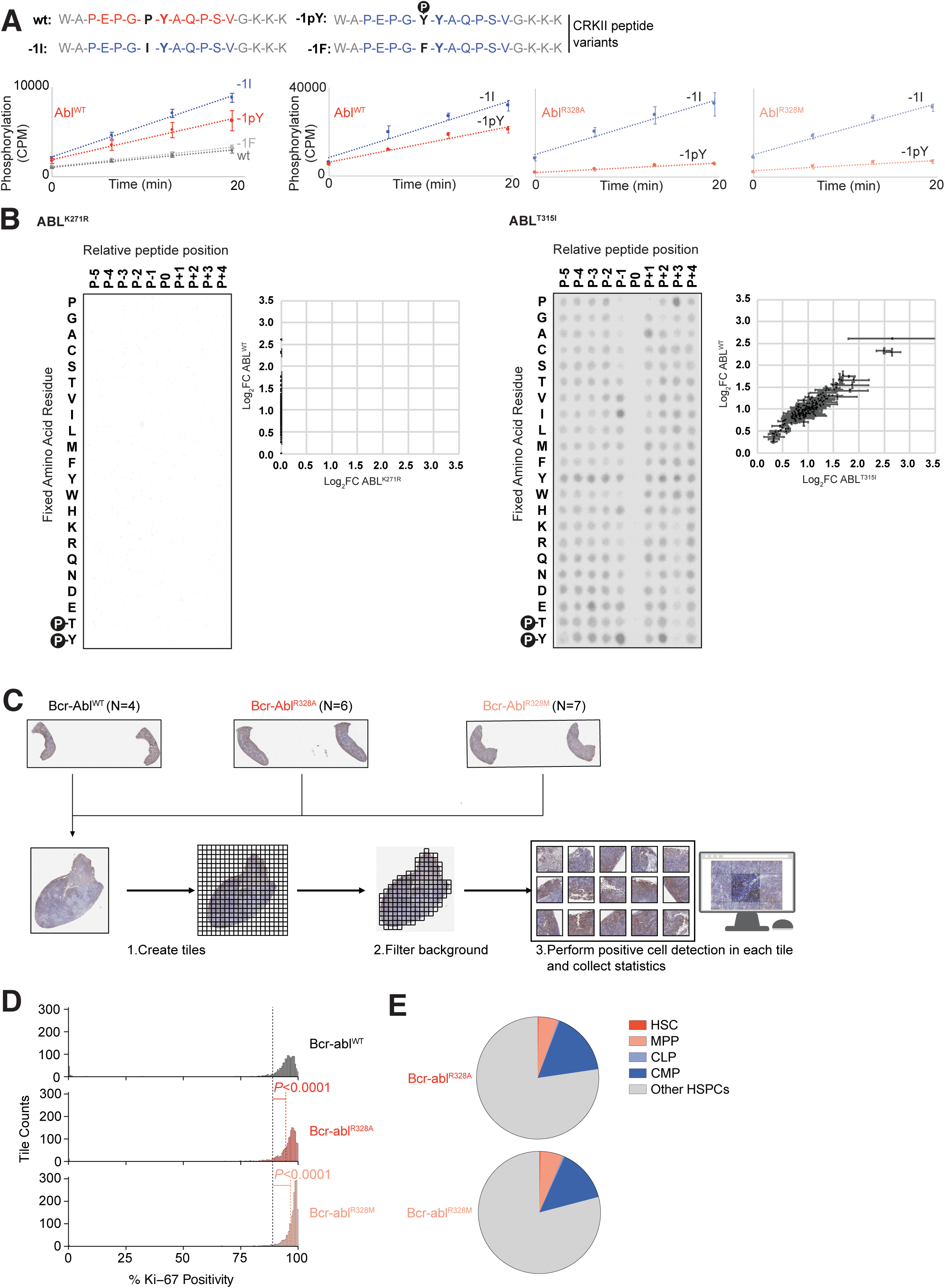
(A) Phosphorylation of specific CrkII peptides (illustrated at the top) by Abl^WT^ (left), Abl^R328A^ (middle) and Abl^R328M^ (right) kinases over time assayed by scintillation counting in triplicate with error bars displaying standard error of the mean. (B) Positional Scanning Peptide Library (PSPL) screening results for the kinase-dead mutant Abl^K271R^ (left) and the hypomorphic mutant Abl^T315I^ (right) kinases with scatterplot quantitatively comparing the column-normalized fold-enrichment values for each spot in the PSPL experiments comparing between them to Abl^WT^ (bottom). Datapoints in the upper-left region of the scatterplot indicate that a specific amino acid residue in a specific position has become less favored or more disfavored, whereas datapoints in the lower-right region of the scatterplot would indicate a change in specificity leading to a more favorable or less disfavorable effect for that amino acid in that position. (C) Workflow for whole-slide image analysis of spleens from mice bearing Bcr-Abl^WT^ (N=4), Bcr-Abl^R328A^ (N=6), or Bcr-Abl^R328M^ (N=7), stained with Ki-67. Slides were tiled, background filtered, and Ki-67 immunopositive cells detected per tile. (D) Full tile-based quantification of Ki-67^+^ cells in spleen IHC sections from mice expressing Bcr-Abl^WT^, Bcr-Abl^R328A^, or Bcr-Abl^R328M^. The complete distribution of Ki-67^+^ tile percentages is shown here, corresponding to the subset ranges (0–5% and 85– 100%) shown in Figure 7H. (E) Pie charts show the distribution of the progenitor cells among HSPC subpopulations from Bcr-Abl^R328A^ and Bcr-Abl^R328M^.

## Supplementary Movies

**Supplementary Movies 1A-C:** Movies showing p27-GFP WT mRuby-PCNA hTert-RPE1 cells treated with either control (A), Abl (B) or Src (C) siRNA. Cells were transfected with siRNA 6 hr before imaging was started.

**Supplementary Movies 2A-C:** Movies showing p27-GFP WT mRuby-PCNA hTert-RPE1 cells treated with either DMSO (A), 10 µM Nilotinib (Abl inhibitor) (B) or 10 µM Saracatinib (C). Cells were treated with inhibitors immediately before imaging was started.

